# Environmental feedback drives oxidative stress response heterogeneity in bacterial populations

**DOI:** 10.1101/2022.09.02.506373

**Authors:** Divya Choudhary, Valentine Lagage, Kevin Foster, Stephan Uphoff

## Abstract

Induction of phenotypic heterogeneity is a ubiquitous consequence of bacterial stress responses. It is commonly postulated that isogenic cells exploit stochastic molecular fluctuations to generate phenotypic heterogeneity as a population survival strategy (termed bet-hedging). However, it is also possible that each cell attempts to maximise its own chances of survival. In that case, the apparent heterogeneity could either be caused by inevitable molecular noise or by underlying deterministic mechanisms which have escaped observation. Here, we investigated the sources and consequences of gene expression heterogeneity in the response of *Escherichia coli* cells to hydrogen peroxide (H_2_O_2_) stress. A machine-learning model accurately predicted the variable responses of individual cells growing in structured populations, indicating that the phenotypic heterogeneity has a deterministic origin. The model further showed that spatio-temporal dynamics in H_2_O_2_ concentration were dictated by the H_2_O_2_ scavenging capacity and morphology of cells in the local environment. Hence, oxidative stress response fluctuations were in fact the result of a precise and rapid feedback between each cell and its immediate environment. Although single cells optimise their individual responses, the formation of short-range H_2_O_2_ gradients by their scavenging activities increases stress tolerance at the population level and leads to spatial variation in mutation rates.

## INTRODUCTION

Bacteria rely on sensory and gene regulatory systems to respond to changes in the conditions of their immediate environment. These systems generate predictable changes in the average gene expression level of cell populations, but the stochastic nature of the underlying molecular processes leads to heterogeneity in the responses between individual cells. Consequently, even cells that experience identical environmental conditions exhibit unpredictable variation in their phenotypes (Spudich & Koshland, 1976). Genes associated with stress responses show particularly high levels of variation (Silander et al., 2012). One explanation is that phenotypic heterogeneity can be beneficial when adverse conditions threaten the survival of a clonal bacterial population (el Meouche et al., 2016; Gefen & Balaban, 2009; Veening, Stewart, et al., 2008; Vincent & Uphoff, 2020); such a bet-hedging strategy can increase the survival probability of the population even though most cells do not maximise their individual fitness (Veening, Smits, et al., 2008). Another explanation for the ubiquity of phenotypic heterogeneity during stress is that gene expression is inherently noisy (Elowitz et al., 2002; Golding et al., 2005; Pedraza & van Oudenaarden, 2005; Rosenfeld et al., 2005; Silander et al., 2012). According to theory, there is a fundamental trade-off between the speed and the accuracy of a gene regulatory response (Lan et al., 2012). When the speed of gene induction is crucial for survival, noise in the response amplitude may be inevitable. For example, oxidative stress caused by exposure to reactive oxygen species (ROS) is rapidly lethal (Barshishat et al., 2018; Imlay, 2013; Ma & Eaton, 1992). Thus, genes of ROS scavenging enzymes (e.g. catalase and peroxidase) are induced in *Escherichia coli* cells within minutes after induction of oxidative stress (Åslund et al., 1999; Lagage et al., 2022). It would thus be expected that these genes exhibit particularly noisy expression, and this was indeed observed in several studies with different types of ROS treatments (de Martino et al., 2016; Lagage et al., 2022; Mitosch et al., 2017; Patange et al., 2018). However, whether phenotypic variation during oxidative stress is truly the result of unavoidable molecular fluctuations, a population bet-hedging strategy, or caused by other factors is unknown.

Besides uncertainty about the functional consequences of phenotypic heterogeneity, its molecular origins are also complex and incompletely understood. Heterogeneity in gene expression at a single-cell level is generally attributed to the stochastic activation and binding of transcription factors(Uphoff et al., 2016), the synthesis of mRNAs and proteins in bursts(Golding et al., 2005), the random partitioning of molecules between sister cells during cell division(Huh & Paulsson, 2011), and changes in gene copy number over the course of DNA replication (M. Wang et al., 2019). These processes lead to variable abundances of proteins even when the average expression rates are constant across cells. Besides instantaneous responses to current conditions, cells also maintain memory of past conditions (Mathis & Ackermann, 2016; Veening, Stewart, et al., 2008; Wolf et al., 2008). The duration of memory is generally set by the cell growth rate, which overall determines the balance between the production and dilution of molecules as cells elongate and divide (Kiviet et al., 2014; Sampaio et al., 2022). Because stress conditions affect growth rates, complex feedback arises between the stress level and gene expression dynamics.

Variation in cell growth and morphology can also affect the influx, dilution, and reaction-diffusion dynamics of stressor molecules such as ROS (Łapińska et al., 2022; Ojkic et al., 2022; Sampaio et al., 2022). Furthermore, cells in a population can interact and modulate their environment in response to stress (Dal Co et al., 2020; Snoussi et al., 2018; van Gestel et al., 2021). As such, it is notoriously difficult to disentangle the primary stochastic sources of phenotypic heterogeneity from secondary deterministic effects. This has been exemplified in studies that revisited the phage lysogenic switch and discovered that its apparent stochastic behaviour can at least partly be explained by previously unobserved deterministic processes of the host cell (Golding, 2018; Snijder & Pelkmans, 2011; St-Pierre & Endy, 2008).

Hydrogen peroxide (H_2_O_2_) is a major ROS that is generated as a by-product of aerobic metabolism under various stresses (Chiang & Schellhorn, 2012; Kohanski et al., 2010; Sen & Imlay, 2021). In addition, bacteria can be exposed to H_2_O_2_ in the environment, for example, from host defences or competing bacterial species (Dong et al., 2015; Imlay, 2019; Passo & Weiss, 1984). Reaction of H_2_O_2_ with iron leads to the formation of hydroxyl radicals that damage DNA and other essential biomolecules (Chiang & Schellhorn, 2012; Gruber et al., 2022; Lagage et al., 2022; Sen & Imlay, 2021). The transcription factor OxyR senses an overabundance of H_2_O_2_ by oxidation and formation of a disulphide bond, and subsequently induces the genes of H_2_O_2_ scavenging enzymes (Åslund et al., 1999; Mishra & Imlay, 2012; Sen & Imlay, 2021). Oxidised OxyR also induces the glutaredoxin GrxA, which reduces OxyR and thereby enables deactivation of the response (Dubbs & Mongkolsuk, 2012). H_2_O_2_ permeates the bacterial cell envelope (Seaver & Imlay, 2001), implying that a higher density of cells that actively scavenge H_2_O_2_ increases the survival of individual cells and by extension the stress tolerance at the population level (Cochran et al., 2000; Ma & Eaton, 1992; Raval et al., 2021).

Overall, these considerations suggest that variability in the responses of bacteria to oxidative stress cannot merely be explained by fundamental gene expression noise but that cellular interactions and growth dynamics could also play an important role. Bacteria may maximise their individual fitness at a single-cell level or exploit phenotypic heterogeneity to improve the chances of population survival. Here, we devised a strategy to pinpoint sources and consequences of oxidative stress response heterogeneity by analysing *E.coli* cells that are growing in structured populations.

## RESULTS

### Micro-scale spatial gradient of the H_2_O_2_ stress response in structured cell populations

We used the “mother machine” microfluidic device for single-cell fluorescence imaging under constant growth conditions (P. Wang et al., 2010). The device consisted of an array of 25 μm long and 1.2 μm wide growth trenches that open to a perpendicular channel with constant medium flow [Figure 1A]. Each trench contained 7 ± 2 cells with an average length of 2.6 ± 0.7 μm per cell (± std) [Figure S1A]. We used a strain with a transcriptional reporter for the OxyR response on a low copy-number plasmid (P*grxA*-CFP) (Zaslaver et al., 2006). A constitutively expressed PRNAI-mKate2 fluorescent protein served as a cytoplasmic label for automated cell segmentation and detection of cell division events. After a period of unperturbed growth, the growth medium flow was changed to medium containing 100 μM H_2_O_2_. Cells activated the *oxyR* response after a delay of 12.4 ± 2.1 min following the onset of treatment. P*grxA*-CFP expression peaked after 38 ± 10.8 min and subsequently stabilised at a lower steady-state expression level that was sustained by the constant flow of fresh medium with H_2_O_2_ into the growth trenches [Figure 1B]. Other transcriptional reporters regulated by OxyR, P*katG*-CFP and *PahpC-CFP*, showed qualitatively similar expression dynamics [Figure S1B-D]. Treatment with 100 μM H_2_O_2_ also caused an instantaneous drop in the rate of cell elongation [Figure 1C]. However, following OxyR activation, cell elongation recovered to pre-treatment rates, indicating complete adaptation to 100 μM H_2_O_2_.

**Figure 1:**
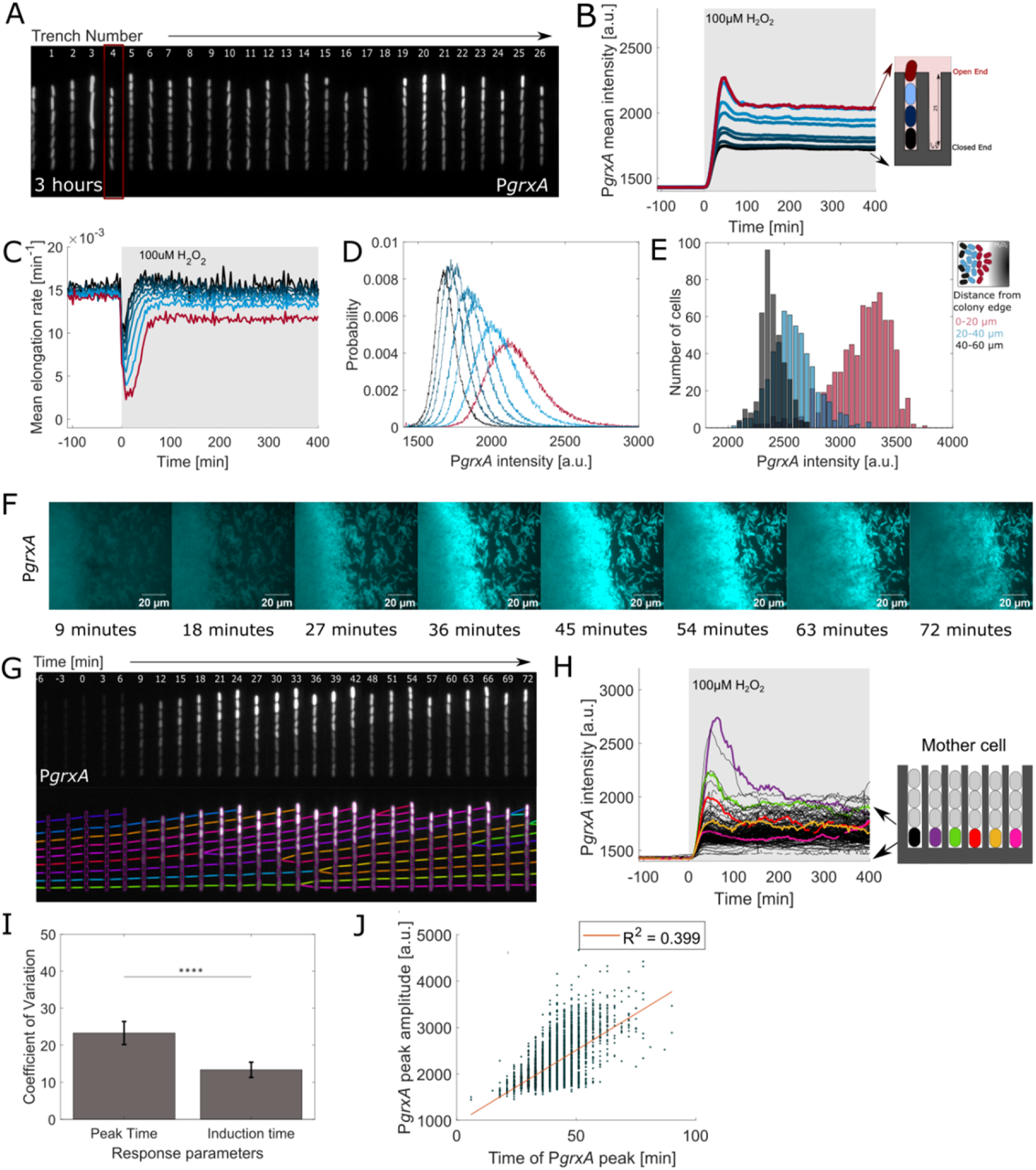
Spatial gradient in oxidative stress response: (A) Snapshot of *E. coli* cells with transcriptional reporter P*grxA*-CFP in growth trenches after 3 hours of 100 μM H_2_O_2_ treatment. (B) P*grxA*-CFP intensities with continuous 100 μM H_2_O_2_ treatment added at time 0 min (shaded area) averaged across cells at specific positions in the growth trench (black line: mother cells at closed end; red line: cells at open end, 3 experimental repeats). (C) Mean log elongation rate for cells at different positions in the growth trench with 100 μM H_2_O_2_ treatment added at 0 min (black line: mother cells at closed end; red line: cells at open end, 3 experimental repeats). (D) Distribution of P*grxA*-CFP intensities of cells with different number of barrier cells computed for 3 to 6 hours after start of 100 μM H_2_O_2_ treatment (black line: mother cells at closed end; red line: cells at open end) (~180,000 data points for steady state histograms, 3 experimental repeats). (E) Histograms of P*grxA*-CFP intensity for cells at different positions in a microcolony after 30 min of 10 mM H_2_O_2_ treatment. (F) P*grxA*-CFP snapshots of a microcolony under 10 mM H_2_O_2_ treatment (Time post treatment annotated in the figures). (G) Kymograph of one growth trench with P*grxA*-CFP intensities and lineage tracing of cells over time with 100 μM H_2_O_2_ treatment added at time 0 min. (H) P*grx*A-CFP intensities of individual mother cells over time (example cells highlighted in colour). (I) Temporal variation of the oxidative stress response across mother cells. Coefficient of Variation (CV, standard deviation/mean) of the response induction time after 100 μM H_2_O_2_ treatment (time until P*grxA*-CFP > 1480 a.u.) and the time to reach the P*grxA*-CFP peak intensity. CV was computed for all mother cells in a single experiment and collated for different repeats. (bar chart represents the mean and standard deviation, **** p<0.0001, 3 experimental repeats). (J) Correlation between P*grxA*-CFP peak amplitude and peak time (each dot represents one mother cell) (~3200 cells, 3 experimental repeats). Orange line represents the linear regression fit with R^2^ =0.399.

The magnitude of the oxidative stress response was highly variable across all cells present in the microfluidic trenches [Figure 1D, S1B]. The mother machine device is popular for studies of cellular heterogeneity because the environmental conditions are believed to be so uniform that any phenotypic variation can be attributed to intracellular noise sources. However, this assumption is rarely tested rigorously. Here, we noticed that cells at the open end of the trenches exhibited higher OxyR reporter expression compared to cells located at the closed end during H_2_O_2_ treatment [Figure 1A-B, D, Movie S1]. By analysing cells according to their position in the trenches, we found that the average magnitude of the OxyR response decreased from the open to the closed end of the trench. The effect of H_2_O_2_ treatment on cell elongation was also reduced with increasing distance from the open end [Figure 1C]. These observations revealed that the oxidative stress response is very sensitive to a cell’s local environment, wherein the stress level decays on a scale of a few micrometres away from the source of treatment. To test the relevance of this steep gradient in bacterial microcolonies, we applied 10 mM H_2_O_2_ treatment (higher concentration than microfluidics to account for cell density dependent protection effect) onto a colony grown on agarose pads and observed a higher *PgrxA*-CFP response along the edge of the colony compared to the interior [Figure 1E-F].

### Machine learning model predicts single-cell responses to H_2_O_2_

Although a cell’s position in a trench accounted for a part of the observed phenotypic heterogeneity, we found that the magnitude of the stress response was variable even between individual cells located at the same position in the trenches, such as the “mother cells’’ positioned at the closed ends [Figure 1G-H]. The magnitude of the initial response peak shortly after treatment showed even higher cell-to-cell variation than the fluctuations of the response during steady-state [Figure S1E]. Notably, the initial delay time of the response was much more uniform across mother cells, whereas the time to reach the expression peak was variable and correlated with the peak response magnitude of each cell (Pearson’s R = 0.399, p = 10^-15^) [Figure 1I-J]. This indicates that while cells can reliably sense the onset of H_2_O_2_ stress and respond rapidly, the response magnitude depends on an unidentified factor that varies substantially between cells.

This heterogeneity could be caused by a variety of mechanisms, including molecular stochasticity in the specific regulatory circuits or general gene expression machinery (Golding et al., 2005; Uphoff et al., 2016), cell-to-cell variation in growth or morphology(Łapińska et al., 2022; Ojkic et al., 2022; Sampaio et al., 2022), variable cell-to-cell interactions (Dal Co et al., 2020; Snoussi et al., 2018; van Gestel et al., 2021), or differences in the local environment of cells (Snoussi et al., 2018; van Vliet et al., 2018). To pinpoint the mechanisms, we designed a machine learning model using random forest regression [Figure 2A]. We computed 126 features based on the time-series data of cell size, shape, and growth rates for each mother cell as well as the other cells present in the same trench (Table S1). The model was trained to predict the peak intensity values of the P*grxA-*CFP reporter after the onset of H_2_O_2_ treatment from 80% of the observed mother cells and tested on the remaining 20% of mother cells (790, unseen during model training). The model predicted the magnitude of the oxidative stress response for individual cells with a mean accuracy of ~70% [Figure 2B, Table S2]. This was surprising as the model did not include any features based on the P*grxA*-CFP signal itself. The apparent noisiness of the stress response is therefore not a consequence of unpredictable molecular fluctuations but has a deterministic origin. Another curious aspect of the model was that it was able to predict P*grxA*-CFP responses for cells treated with a range of H_2_O_2_ concentrations (37.5 – 100 μM) although the concentration itself was unknown to the model. Hence, the features extracted from cell growth characteristics and morphology were sufficient to deduce the underlying H_2_O_2_ concentration [Figure S2B, Table S3, S4].

**Figure 2:**
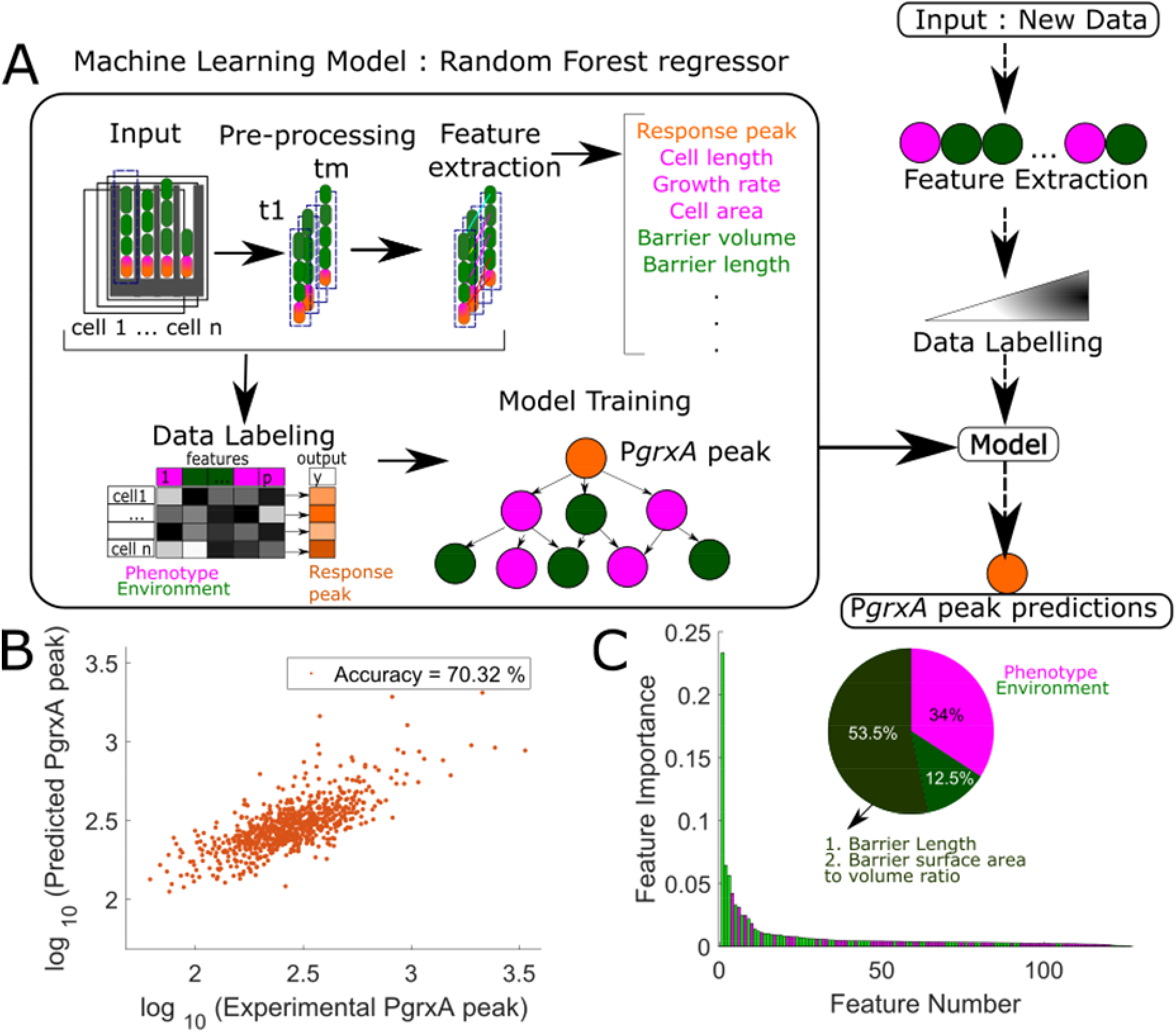
Machine learning predicts single cell response heterogeneity: (A) A random forest machine learning model predicts P*grxA*-CFP peak intensities of ~789 mother cells (orange). It uses features that describe the phenotypic characteristics of the mother cell (magenta) and the other cells in the local environment of each trench (barrier cells, green). (B) P*grxA*-CFP peak predicted by the model plotted against the experimentally measured P*grxA*-CFP peak (each dot represents one mother cell) (data in Table S2). (C) Feature importance plots show the relative contribution of the 126 input features to the predictive power of the model (feature names in Table S1). Barrier cell features (green) are more important than the mother cell features (pink). The two most important features are highlighted and account for 53.5% of the predictive power.

Our machine learning approach eased the way to pinpoint the sources of cellular heterogeneity. The features that contributed the most to the predictive power (~66%) of the model related to the characteristics of what we term “barrier cells”, i.e. the cells located between the mother cell and the open end of each trench [Figure 2C]. The strongest features accounting for ~53.5% of the total predictive power were (i) the cumulative length of the barrier cells in a trench, and (ii) the combined surface area-to-volume ratio of all the barrier cells in a trench. In fact, a model that contained only features of the barrier cells and no features of the mother cell was still able to predict mother cell responses with high accuracy [Figure S2A]. Hence, it appears that an individual cell’s stress response magnitude does not depend on its own characteristics or location but can be accurately predicted from the number and morphology of its neighbours.

### Variation in the spatial structure of the micro-environment causes oxidative stress response heterogeneity

To understand the apparent role of cell-to-cell interactions in the oxidative stress response, we designed a microfluidic experiment with a variable number of cells per trench at the time of treatment (5 ± 2 cells) [Figure S3A], instead of filling all growth trenches completely. Under these conditions, the magnitude of the oxidative stress response of the mother cells became even more variable [Figure S3B-C]. P*grxA*-CFP intensities of mother cells decreased when an increasing number of barrier cells was present, suggesting that these are protecting the mother cell from H_2_O_2_ [Figure 3A-B]. The number of barrier cells fluctuates as cells divide and are pushed out of the open end of the trenches. Although the cell-average reporter expression was stable during constant H_2_O_2_ treatment, individual mother cells showed dynamic fluctuations at steady-state [Figure 3C]. Cross-correlation analysis showed that these expression fluctuations were negatively correlated with the variation in the number of barrier cells in the same trench [Figure 3D, S4]. The negative cross-correlation peaked at a lag time of 4.5 ± 1.5 min, showing that fluctuations in the number of barrier cells preceded changes in P*grxA*-CFP expression. Hence, the OxyR response is exquisitely sensitive and rapidly responds to very subtle changes in a cell’s micro-environment, such as the division or disappearance of a single cell in the vicinity. Based on calibration experiments with different H_2_O_2_ concentrations [Figure S5A-B], we inferred the local extracellular H_2_O_2_ concentration from the P*grxA*-CFP intensity of a cell. This showed an exponential decrease of H_2_O_2_ concentration from the open end of the trench and each barrier cell decreased the local H_2_O_2_ concentration by 32.3 ± 4.6% [Figure S5C].

**Figure 3:**
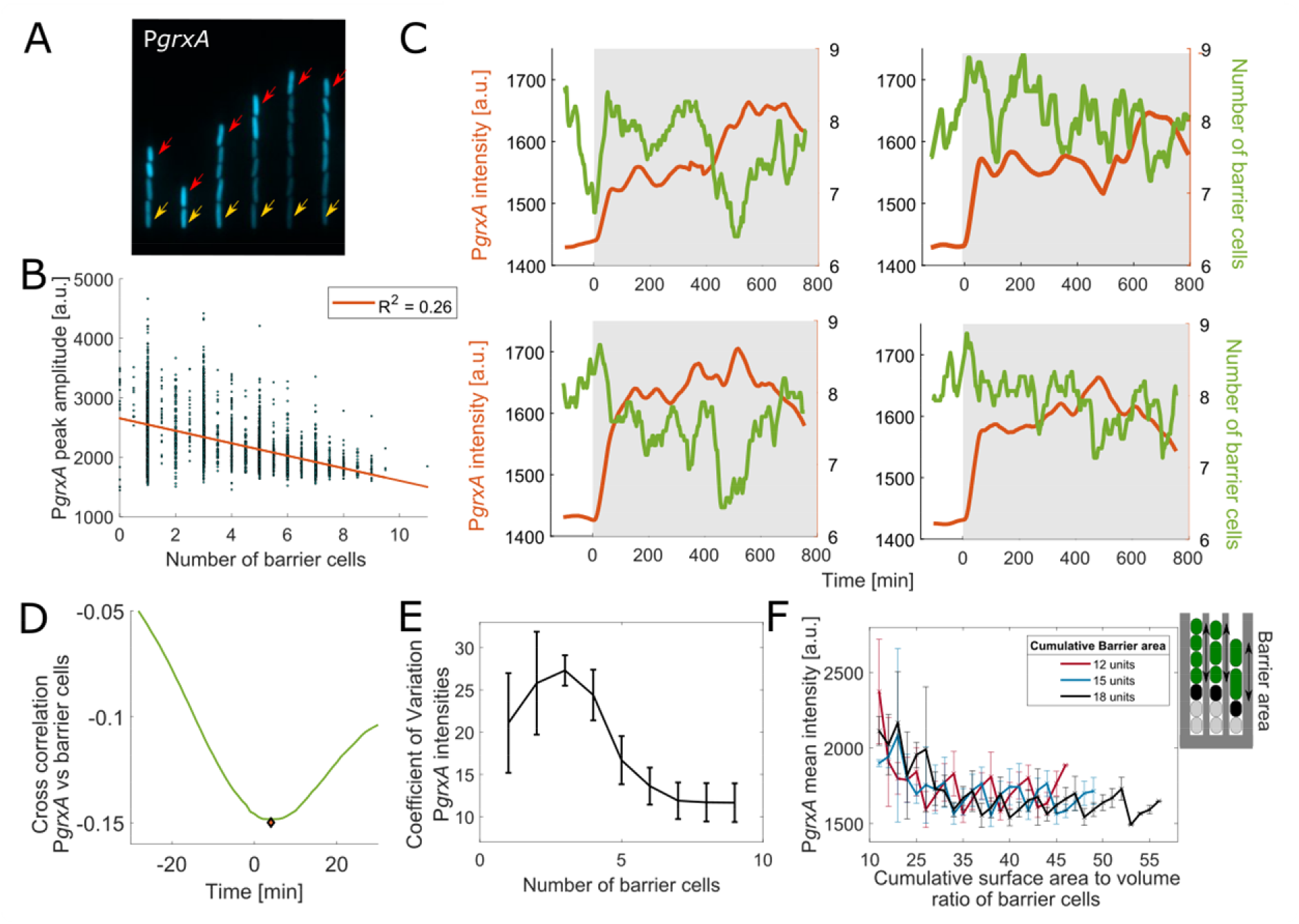
Variation in the spatial structure of the micro-environment explains response fluctuations: (A) Snapshot of P*grxA*-CFP under 100 μM H_2_O_2_ treatment for growth trenches with variable number of cells (mother cells marked with yellow arrow, outermost cells with red arrow). (B) P*grxA*-CFP peak intensity amplitude under 100 μM H_2_O_2_ treatment plotted against the average number of barrier cells around the time of treatment (± 9 min; each dot represents one mother cell) (~3250 cells, 3 experimental repeats). (C) Example time-traces of P*grxA*-CFP intensity (orange) for mother cells and the number of barrier cells in the same trench (green) with 100 μM H_2_O_2_ treatment added at 0 min (shaded area). Curves were smoothed using a moving mean filter with 45 min window. (D) Mean temporal cross correlation for P*grxA*-CFP of mother cells against the number of barrier cells per trench (example time traces shown in panel C), when mean P*grxA*-CFP intensity has reached steady-state from 2 hours after start of 100 μM H_2_O_2_ treatment until end of experiment (~11 hours) (~950 cells, 2 experimental repeats). Minima is represented by the orange diamond. (E) Coefficient of Variation for P*grxA*-CFP intensity of mother cells with different number of barrier cells (CV: std/mean was computed for all mother cells in a single experiment, error bars represent std across experiments). (F) P*grxA*-CFP intensity for cells with the same total area of barrier cells (18, 15 or 12 units) plotted against the cumulative surface area to volume ratio of the barrier cells (3 experimental repeats).

The response variation persisted even when restricting the analysis to mother cells with a fixed number of barrier cells, especially when that number was small [Figure 3E]. This indicates that variation in the characteristics of the barrier cells also propagates to the mother cell, whereas such variation averages out when the number of barrier cells is larger. Indeed, the Machine Learning model pointed at the surface area-to-volume ratio of the barrier cells as an important feature that determines mother cell responses besides the total length of the barrier cells. To test this directly, we reasoned that an increased number of barrier cells at a constant cumulative volume would result in a larger total surface area-to-volume ratio. As predicted, we found that the P*grxA*-CFP magnitude decreased with the surface area-to-volume ratio of barrier cells while their cumulative volume was kept constant [Figure 3F]. Therefore, fluctuations in the number, size, and morphology of neighbouring cells define an individual cell’s dynamic response to H_2_O_2_. Notably, the basal expression of P*grxA*-CFP without H_2_O_2_ treatment was more uniform across cells and unaffected by any variation in barrier cells [Figure S5D].

### Intracellular scavenging enzymes create a local H_2_O_2_ gradient

To explore how barrier cells create an exponential H_2_O_2_ gradient, we used microfluidic chips in which the trenches were only partially filled with cells. The outermost cells in each growth trench exhibited the same level of stress response irrespective of their distance from the open end, showing that the geometry of the trenches itself does not restrict H_2_O_2_ diffusion [Figure 3A, 4A]. However, widening the growth trenches from 1.2 μm to 1.4 μm reduced both the barrier effect [Figure 4B] and cell-to-cell variation [Figure S6], showing that a decrease in cell density permits a higher and more uniform H_2_O_2_ flux. In principle, this effect could be caused by the cell mass passively blocking H_2_O_2_ diffusion, or by active degradation of H_2_O_2_ by the barrier cells.

**Figure 4:**
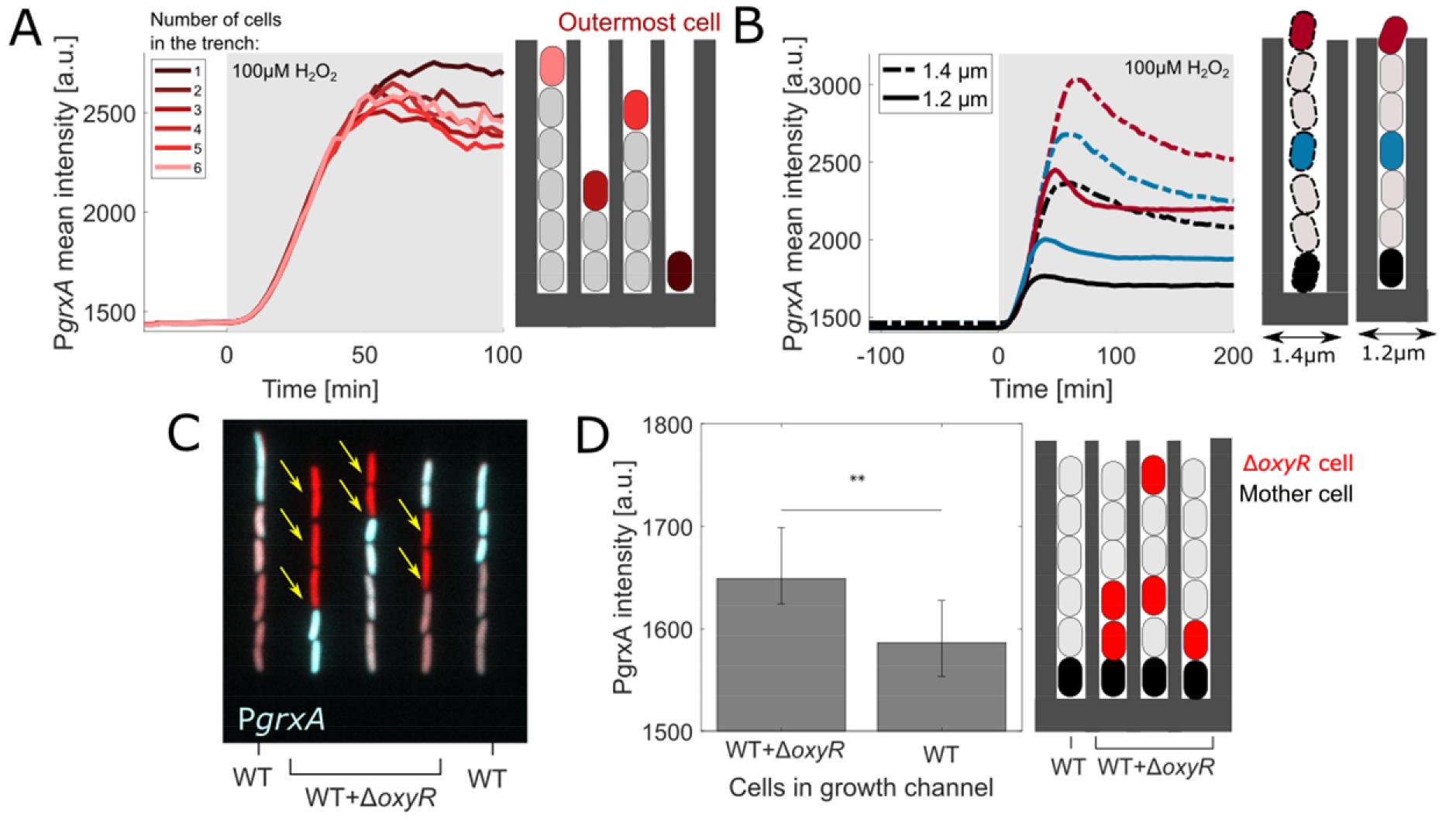
Intracellular scavenging enzymes create short-range local H_2_O_2_ gradients: (A) Mean P*grxA*-CFP intensity for outermost cells in growth trenches (marked by red arrows in Figure 3A with total number of cells per trench ranging from 1 to 6). 100 μM H_2_O_2_ treatment added at 0 min (shaded area) (4 experimental repeats). (B) P*grxA*-CFP mean intensities for trenches with widths of 1.2 μm (solid lines) and 1.4 μm (dashed lines) for cells with 0 (red), 3 (blue) and 6 (black) barrier cells (3 experimental repeats each). (C) Snapshot of merged P*grxA*-CFP (cyan) and mKate2 cell marker (red) intensity for trenches with a mix of WT and Δ*oxyR* strain under 100 μM H_2_O_2_ treatment (Δ*oxyR* cells marked with arrow). (D) Mean P*grxA*-CFP intensity for WT mother cells in growth trenches containing a mix of barrier cells with at least one Δ*oxyR* cell or purely WT barrier cells (~1600 mother cells, 3 experimental repeats, error bars represent 25^th^ and 75^th^ percentile, ** p<0.01).

The latter mechanism is likely considering that H_2_O_2_ is rapidly scavenged by catalase which is induced by the OxyR response. The Damköhler number (a dimensionless quantity that relates the reaction rate to the rate of diffusion) is ~4·10^3^ for the ratio of the catalase scavenging rate (5.4·10^4^ s^-1^, (Mishra & Imlay, 2012)) to the diffusion of H_2_O_2_ across the *E. coli* cell envelope (1.6·10^-5^ m/s, (Seaver & Imlay, 2001)). Hence, the scavenging rate is limited by the diffusion into cells, coherent with our finding that the protective effect of the barrier cells was determined to a large extent by their surface area. To experimentally verify the contribution of H_2_O_2_ scavenging towards the creation of an exponential stress gradient, we loaded trenches with a mix of a Δ*oxyR* strain and a wild-type strain with P*grxA*-CFP reporter [Figure 4D]. Δ*oxyR* cells can scavenge endogenously produced H_2_O_2_ but cannot induce catalase and other oxidative stress response genes. Indeed, the gradient in P*grxA*-CFP reporter intensity was disrupted and wild-type cells positioned behind Δ*oxyR* cells showed elevated stress responses [Figure 4C-D, S8A, Movie S2]. In contrast, wild-type cells had a marked protective effect on Δ*oxyR* cells, consistent with bulk culture measurements (Ma & Eaton, 1992). On their own, Δ*oxyR* cells were unable to grow during 100 μM H_2_O_2_ treatment [Figure S7, Movie S3]. The presence of wild-type cells acting as a barrier rescued the growth of Δ*oxyR* cells until the wild-type cells exited the trench [Movie S4-5]. Bacterial colonies where wild-type strain with P*grxA*-CFP reporter was mixed with the Δ*oxyR* strain exhibited a higher and more uniform stress response from the edge to the interior than in a clonal wild-type colony [Figure S8B-D].

### Barrier cells obtain high H_2_O_2_ tolerance via gradual adaptation

Variation in the local H_2_O_2_ concentration determined divergent cell fates after H_2_O_2_ treatment [Figure 5A]. A 1-hour pulse with 500 μM H_2_O_2_ concentration stalled the growth of 99% of mother cells [Figure 5B]. We found that 13% of mother cells subsequently recovered growth and these individuals were protected by more barrier cells than those that irreversibly stopped growth [Figure 5B]. Furthermore, the delay before the surviving cells started to regrow after treatment removal was variable and negatively correlated with the number of barrier cells [Figure 5A, C]. Elongation rates of the outermost cells in the trenches revealed that >50 μM H_2_O_2_ led to growth inhibition in the absence of cellular cross-protection [Figure S9]. The onset of 100 μM H_2_O_2_ treatment completely halted the growth and division of the ~1-2 outermost barrier cells per trench, while the cells in the interior of the trenches only transiently slowed in growth and quickly recovered unperturbed elongation and division rates despite ongoing H_2_O_2_ treatment [Figure 1C]. This raises the question of how the barrier is first replenished and then able to withstand apparently lethal concentrations of H_2_O_2_ to provide continuous protection for the cells in the interior. After the initial demise of the outermost cells, the surviving interior cells divide and push live progeny towards the stress source and thereby replace the barrier cells that had been killed by the sudden onset of treatment. Therefore, rather than being exposed suddenly, barrier cells are able to adapt to a gradual increase in H_2_O_2_ concentration. Such gradual adaptation is known to enhance survival of higher stress levels (Imlay, 2008; Lagage et al., 2022; Rodríguez-Rojas et al., 2020). To confirm this “priming” effect underlies the high stress tolerance of the barrier cells, we applied a gradual increase of H_2_O_2_ concentration reaching up to 500 μM over 18 minutes [Figure 5A], which greatly increased the fraction of cells that recovered growth after H_2_O_2_ removal, compared to sudden treatment with 500 μM H_2_O_2_ [Figure 5B].

**Figure 5:**
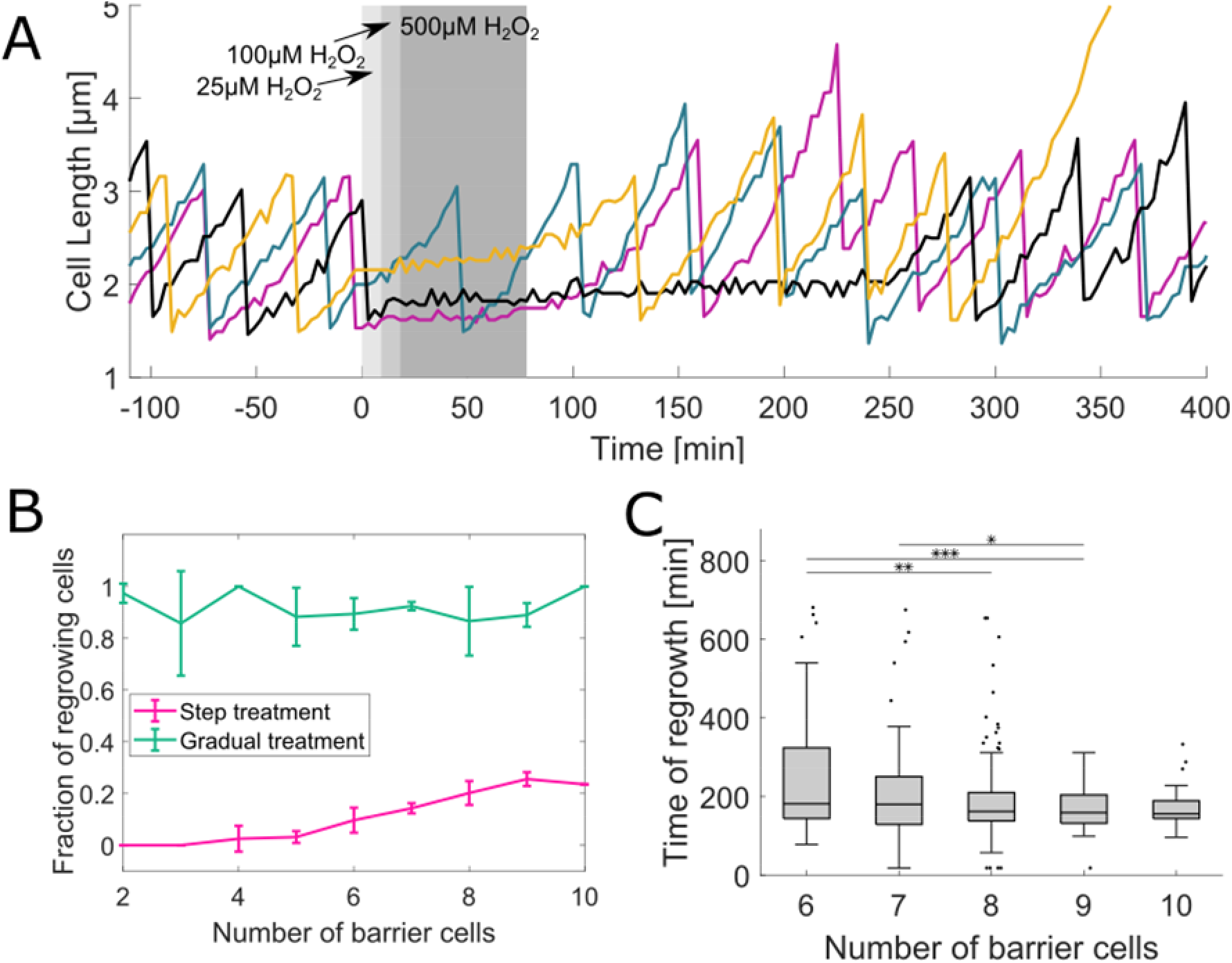
Local environmental variation determines divergent cell fates after H_2_O_2_ treatment: (A) Individual mother cell length traces with gradual increase in H_2_O_2_ concentration from 25 μM to 100 μM to 500 μM (shaded area), followed by recovery without treatment. (B) Proportion of mother cells that recover growth within ~11 hours after the removal of 500 μM H_2_O_2_ treatment that was applied in a single step (pink) or gradually (green, as illustrated in panel A) plotted against number of barrier cells at the time of treatment (≥3 experimental repeats each; line plot with mean and standard deviation). (C) Time of regrowth of mother cells after gradual 500 μM H_2_O_2_ treatment (as illustrated in panel A) plotted against number of barrier cells at the time of treatment. (~300 cells; boxplot with median, 25^th^ and 75^th^ percentile; *p<0.05, ** p<0.01, *** p<0.001).

To understand the spatio-temporal dynamics of adaptation, we tracked individual cells as they moved from birth at the closed end of the trench until they exited the open end [Figure 6A]. P*gxr4*-CFP expression increased rapidly over time as cells traverse the spatial H_2_O_2_ gradient [Figure 6B, S10]. This is caused by two effects; first, the exponential increase in H_2_O_2_ concentration along the length of the trench, and second, the increasing speed of movement as cells are pushed by exponentially growing cells located deeper in the trench. The speed varies between cells due to fluctuations in elongation rates and the number of cells per trench [Figure 6C]. Cells that had the same number of barrier cells and moved faster to the trench opening showed a steeper response than slower moving cells [Figure 6D]. This effect was not seen in untreated conditions [Figure S11]. When mobile cells experience a gradient of H_2_O_2_ in space and time, successful adaptation requires that the induction rate of scavenging enzymes matches the increasing influx of H_2_O_2_. Indeed, movement speed and the response induction rate were correlated in individual cells (Pearson correlation coefficients = 0.76) [Figure 6E]. Next, in order to isolate the effect of the spatial gradient on the response induction rate, we analysed cells that moved with similar speed. In this case, cells with fewer barrier cells had a higher response induction rate because they experience a steeper spatial H_2_O_2_ gradient [Figure 6F]. Together, the multiplicative effects of accelerated movement and increasing stress gradient lead to a rapid induction of the response as cells are pushed towards the trench opening.

**Figure 6:**
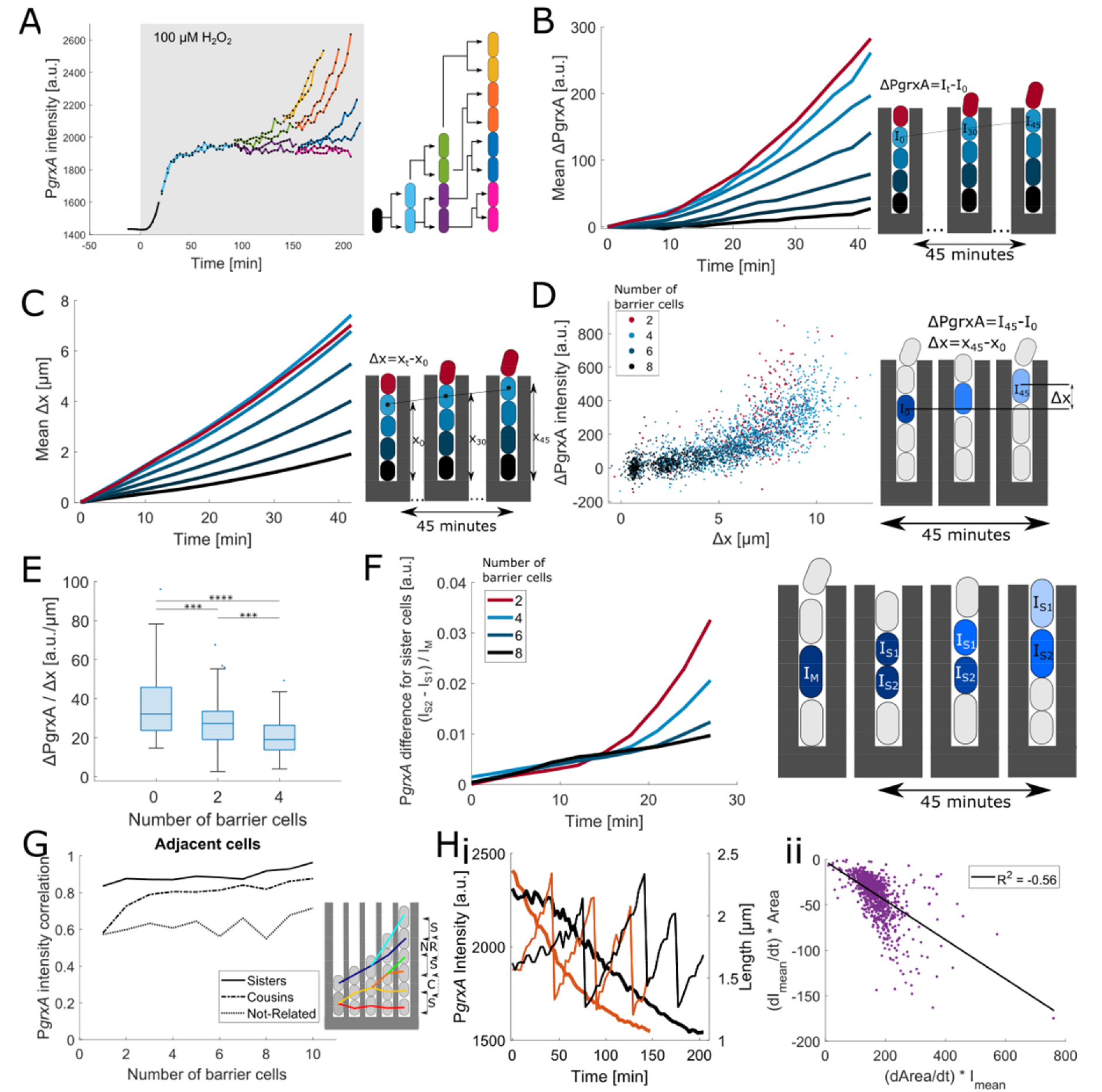
Spatio-temporal response dynamics overwrite cell lineage-associated memory: (A) P*grxA*-CFP intensities for a representative mother cell and its progeny that move towards the opening of the growth trench. 100 μM H_2_O_2_ treatment added at 0 min (shaded area). (B) Effect of cell movement on P*grxA*-CFP expression in a spatial H_2_O_2_ gradient: Mean increase in P*grxA*-CFP intensity (ΔPgrxA) over time when the population response has reached steady-state (from 2 hours after start of 100 μM H_2_O_2_ treatment until end of experiment ~11 hours). Each curve represents cells moving towards the trench opening from a different starting position (black line: mother cells at closed end; red line: cells at open end) (3 experimental repeats). (C) Effect of cell position in the growth trench on the movement speed: Each curve shows the mean movement Δx towards the trench opening of cells with different starting position (black line: mother cells at closed end; red line: cells at open end) (3 experimental repeats). (D) Response induction rate depends on the cell movement speed: Each point represents the total increase in P*grxA*-CFP and total movement of a single cell traced for 45 min. Colours represent the starting position according to the number of barrier cells (~2800 total number of cells, 3 experimental repeats). (E) Response induction rate depends on cell position in the growth trench: Relative increase in P*grxA*-CFP per distance for cells that move 7 ± 0.2 μm in 45 min. Number of barrier cells represents the starting position (~200 total number of cells, 3 experimental repeat, *** p<0.001, **** p<0.0001). (F) Responses of sister cells diverge over time: P*grxA*-CFP intensity difference between sister cells (S1 and S2) followed for 30 minutes post division when the response has reached steady-state. Intensity differences were normalised by the intensity of the cell they arose from. Number of barrier cells represents the starting position at the time of cell division (~5400 pairs, 3 experimental repeats). (G) Effect of cell position on response memory: Pearson’s correlation coefficient for P*grxA*-CFP intensities of adjacent cell pairs with variable number of barrier cells, related as sisters (S, solid), cousins (C, dash-dotted) or not related (NR, dotted) (3 experimental repeats). (H) Heterogeneous growth rates explain variability in the deactivation rate of the response: (i) P*grxA*-CFP intensity (thick lines) and cell lengths (thin lines) of 2 representative mother cells (black and orange plots respectively) after removal of 500 μM H_2_O_2_ at time 0 min, showing that faster growth leads to a faster decay. (ii) Correlation between PgrxA-CFP intensity decay (dI_mean_/dt) and cell growth rate (dArea/dt) after removal of 500 μM H_2_O_2_ (~1000 mother cells).

### Rapid response to environmental fluctuations overwrites cellular memory

In principle, a cell’s gene expression level is shaped by its immediate response to the current local environment as well as its memory of past stress exposure (Mathis & Ackermann, 2016). We thus explored if response heterogeneity can be traced back to a cell’s lineage history. Cytoplasmic proteins are randomly equipartitioned at cell division, so the CFP reporter intensities of sister cells were closely correlated immediately after birth [Figure S12A]. However, the intensity of the cell located closer to the open end of the trench quickly diverged from the sister [Figure 6F, S12B]. Specifically, the intensity difference between the two sisters increased exponentially over time because the cell closer to the open end of the trench experiences a higher H_2_O_2_ concentration and itself acts as a barrier for the cell located below. This effect was much less pronounced for sisters born with a larger number of barrier cells because they move more slowly in a shallower H_2_O_2_ gradient. Hence, any stress response memory is erased quickly for cells that move rapidly up the H_2_O_2_ gradient at the open end of the trenches, whereas cells at the closed end experience more uniform conditions and could therefore maintain memory across generations.

To test this, we quantified the difference in CFP reporter intensities between adjacent cells that were either related as sisters (separated 1 generation before), cousins (2 generations separated), or not related (>2 generation separated). There was less variation in the responses between sister pairs than between cousins or distantly related cell pairs [Figure 6G, S13A]. As predicted, sisters and cousins became more correlated with increasing number of barrier cells [Figure 6G, Figure S13B]. Therefore, although oxidative stress response memory can in principle persist over generations when the environment is constant, changing environmental conditions dictate the instantaneous response level and effectively overwrite cell lineage memory. When H_2_O_2_ treatment was stopped, reporter intensities started to drop quickly and decayed exponentially with a half-life that matched the cell doubling time [Figure S14A-B]. The P*grxA*-CFP decay rate was variable across cells but closely correlated with the growth rate, which explained the observed cell-to-cell variation in the response deactivation [Figure 6H].

### Spatial gradients in H_2_O_2_ lead to cellular heterogeneity in mutagenesis

Because ROS can generate mutagenic DNA damage, we reasoned that spatial and temporal fluctuations in ROS concentrations may lead to mutational heterogeneity across cells in a population. It has been shown that variation in the timing (Uphoff, 2016; Vincent & Uphoff, 2021) or magnitude of a stress response (Pribis et al., 2019) can cause cell-to-cell variation in mutation rates, but potential effects of spatial stress gradients on mutagenesis have not been explored. We previously reported that H_2_O_2_ treatment causes a burst of mutations prior to induction of the oxidative stress response in *E. coli* (Lagage et al., 2022). Here, we used the same approach with a MutL-mYPet fusion (which forms foci at sites of DNA mismatches) to monitor the occurrence of DNA replication errors in single cells (Robert et al., 2018; Uphoff, 2018). Cells showed a burst of DNA replication errors immediately after the onset of treatment [Figure 7A-B], and the magnitude of this burst decreased steeply with increasing number of barrier cells [Figure 7C]. There was no spatial gradient in mutagenesis for cells growing in wider trenches [Figure 7C], consistent with the uniform stress response observed under these conditions. Furthermore, bacterial colonies with 10 mM H_2_O_2_ treatment exhibited a higher frequency of DNA mismatch foci along the edge of the colony compared to the interior [Movie S6]. Therefore, mutational heterogeneity can arise from local cellular interactions that generate spatial variation in stress level between cells.

**Figure 7:**
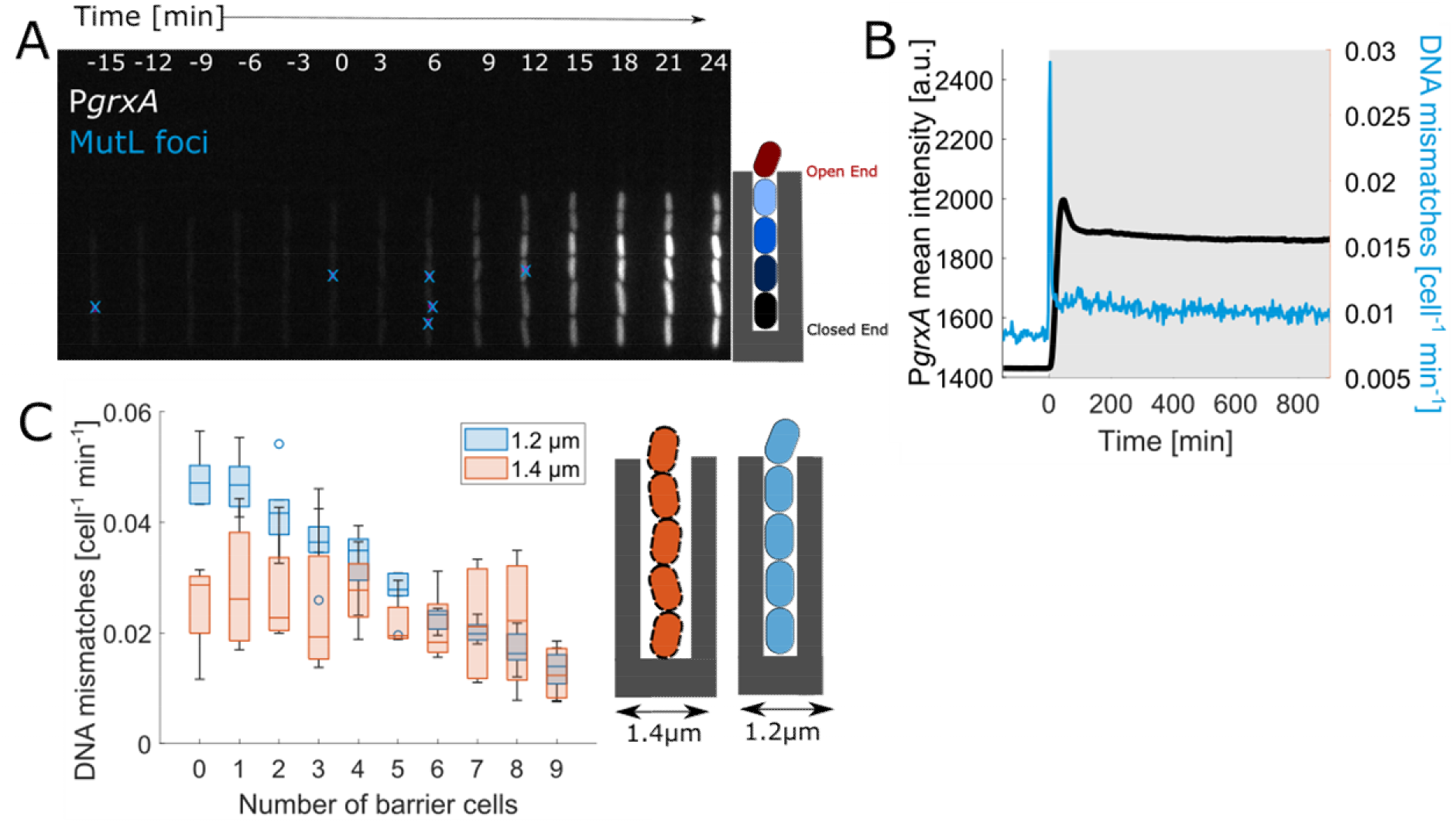
Spatial heterogeneity in H_2_O_2_ concentration causes cell-to-cell variation in mutagenesis: (A) Kymograph of one growth trench with P*grxA*-CFP intensities under 100 μM H_2_O_2_ treatment added at time 0 min. Blue crosses represent MutL-mYPet mismatch foci. (B) Mean rate of DNA mismatch foci (per cell per minute, blue) and P*grxA*-CFP mean intensity (black) for all cells in the growth trenches (3 experimental repeats). (C) Dependence of the mismatch rate on cell position: Amplitude of the DNA mismatch rate peak for cells with different number of barrier cells under 100 μM H_2_O_2_ treatment for growth in 1.2 μm (blue) and 1.4 μm (orange) wide trenches (3 experimental repeats each).

## DISCUSSION

The presence of phenotypic heterogeneity within bacterial populations has been well documented. However, the exact sources and functional consequences of the heterogeneity are generally still difficult to pinpoint. One can distinguish three possible scenarios: (i) The heterogeneity serves a functional purpose for the fitness of the population. Bacteria can utilise stochastic molecular processes to randomise their behaviour as a bet-hedging strategy, such that at least a fraction of cells is well adapted to unpredictable environmental changes. (ii) The heterogeneity does not have any functional benefit itself but reflects the limited accuracy by which bacteria can sense their environment and regulate responses due to unavoidable biochemical noise. For example, bacteria may prioritise the speed of a response at the cost of accuracy. (iii) Individual cells maximise their own fitness accurately and the population heterogeneity is driven by an underlying deterministic process that remains hidden or appears random because it has not been thoroughly characterised yet. These scenarios are not necessarily exclusive, and a part of the apparent heterogeneity may also stem from non-biological measurement noise.

Studies on the oxidative stress response in bacteria have revealed characteristics that partially support each of these contrasting models (Cochran et al., 2000; de Martino et al., 2016; Hong et al., 2020; Patange et al., 2018; Sampaio et al., 2022). To resolve this uncertainty, we characterized the spatio-temporal behaviour of single *E. coli* cells growing in defined structured populations under a constant treatment of H_2_O_2_. We observed micrometerscale spatial gradients of the stress response across cells in the population. Furthermore, individual cells at fixed positions in the population exhibited substantial fluctuations in gene expression. Machine learning analysis complemented with microfluidic experiments not only showed that the response fluctuations had a largely deterministic origin but the approach also eased the screening of a large number of features of single cells and their micro-environment to understand the underlying mechanism. Contrary to the apparent noisiness of the response, individual cells in fact tune their gene expression extremely precisely to the local H_2_O_2_ concentration. Induction of scavenging enzymes generates steep spatial H_2_O_2_ gradients, with each cell contributing to a ~30% reduction of H_2_O_2_ in its vicinity. Temporal variation in H_2_O_2_ concentrations result from subtle changes in the scavenging capacity and surface area of cells. Ultimately, seemingly random fluctuations in stress response expression are dictated by tight feedback between a cell and its local environment.

The accuracy of the oxidative stress response is likely based on the rapid uptake of H_2_O_2_ together with its high reactivity and specificity towards OxyR (Lee et al., 2004). Induction of scavenging enzymes and GrxA reductase lowers the intracellular H_2_O_2_ concentration and deactivates OxyR in a negative feedback loop, which increases response speed and accuracy in general (Alon, 2007). The formation of short-range ROS gradients protects bacteria residing in communities such as biofilms, and may also reflect the potential utility of these molecules for local cell-to-cell communication (D’Autréaux & Toledano, 2007; Erttmann & Gekara, 2019). Because oxidative stress is mutagenic, gradients in H_2_O_2_ concentration cause spatial variation in the frequency of DNA replication errors across cells. Therefore, local environmental heterogeneity can diversify mutation rates among cells growing in a structured population. Indeed, our observation that H_2_O_2_-induced mutation rates decrease with increasing cell density matches results obtained in liquid bulk culture (Krašovec et al., 2014, 2017).

Although our results overall support scenario (iii) that individual cells optimise their own oxidative stress response according to the ROS levels in their immediate environment, this strategy also leads to benefits at the population level as predicted by scenario (i). Sudden exposure to high H_2_O_2_ concentrations kills cells on the outside of a dense population, but the initial H_2_O_2_ gradient created by these cells allows the survivors in the interior to regrow and replenish the gradient for continuous protection. Therefore, the spatial gradient also has a temporal effect by allowing cells to traverse a gradual increase in stress level and obtain high tolerance via the priming effect.

We expect our combination of single-cell imaging and machine-learning to be generally applicable towards understanding sources of heterogeneity and its effects on stress adaptation from single cells to populations.

## ACKNOWLEDGEMENTS

We thank Helen Alexander, David Sherratt, Mathew Stracy and members of the Uphoff lab for their discussions and comments on the manuscript. Research in the Uphoff lab is funded by a Wellcome Trust & Royal Society Sir Henry Dale Fellowship (206159/Z/17/Z), a Wellcome-Beit Prize (206159/Z/17/B) and a Research Prize Fellowship of the Lister Institute of Preventative Medicine. D.C was supported by an Oxford-Indira Gandhi Scholarship funded by the Oxford India Center for Sustainable development.

## AUTHOR CONTRIBUTIONS

Conception and design of study: DC, VL, KF and SU, Engineering of genetic constructs: DC, VL and SU, Data collection: DC, Development of statistical analysis: DC and SU, Data analysis and interpretation: DC and SU, Writing of the article: DC and SU.

