## Supplementary material for "Environmental feedback drives oxidative stress response heterogeneity in bacterial populations": Materials and methods and supplementary information

#### Strains and plasmids

Experiments were performed with strains derived from *E. coli* K12 AB1157. The strains constitutively expressed $P_{RNAI}$-mKate2 fluorescent marker for cell segmentation analysis and the *flhD* gene was deleted to inhibit flagellar motility allowing growth in microfluidic chips. This strain also carried an endogenous MutL-mYPet fusion for detection of DNA mismatches (Uphoff, 2018). The strain SU802 with a Δ*oxyR*::kan deletion was described in (Lagage et al., 2022). The strain 882 with a Δ*oxyR*::kan deletion also carried a pUC18 plasmid expressing YPet as a fluorescent marker to distinguish Δ*oxyR* cells from wild-type cells in the experiments with a mixture of the two strains.

The OxyR response reporter plasmids (P*grxA*-CFP, P*katG*-CFP and P*ahpC*-CFP) were derived from an *E. coli* promoter library of pSC101 plasmids (Zaslaver et al., 2006). Each plasmid in the library contains the promoter region of a specific gene or operon in front of GFPmut2 fluorescent protein. To avoid overlap with the yellow fluorescence channel for imaging MutL-mYPet, we changed the GFPmut2 to the fast-maturing cyan fluorescent protein SCFP3A using Gibson Assembly (NEB).The promoter regions in the reporter plasmids were sequenced and plasmids were transformed into the strain SU178. Presence of the expected fluorescent protein signals were verified by taking microscopy snapshots. Strains were selected for and subsequently grown in LB, or LB agarose supplemented with 25 µg/mL kanamycin.

#### Media and growth conditions

Strain construction was performed in LB or LB supplemented with antibiotics at 37˚C. 4 mL cultures were grown at 200 rpm in 10 mL culture tubes. Successfully constructed strains were stored in glycerol stocks at -80˚C. For experiments, strains were streaked from these glycerol stocks on LB agarose plates with appropriate antibiotic selection. A single colony was picked and then grown overnight in M9 minimal media. This media was prepared with M9 salts (15 g/L $KH_2PO_4$, 64 g/L $Na_2HPO_4$, 2.5 g/L NaCl, and 5.0 g/L $NH_4Cl$), 2 mM $MgSO_4$, 0.1 mM $CaCl_2$, 0.5 mg/mL thiamine, MEM amino acids, 0.1 mg/mL L-proline, and 0.2% glucose. The next day, overnight culture was diluted 1:50 and grown to OD600 ~0.3 in M9 minimal media. For loading cells in microfluidic chips, 0.85 mg/mL Pluronic F127 was added to the media to avoid cell aggregation. For experiments done under hydrogen peroxide treatment, the specific concentration of $H_2O_2$ was added to the growth media immediately before the start of the experiment.

#### Microfluidic chip preparation

Single-cell imaging was performed using the 'mother machine' microfluidic device as described in (Lagage et al., 2022; Wang et al., 2010). The chip has a main channel for flow of media with dimensions 25 µm width and 100 µm height. This main channel is branching into perpendicular growth channels (here called 'growth trenches') of dimension 1.2 µm width and 1.2 µm height and 25 µm length. Where indicated, a different silicon wafer was used with larger trenches of 1.4 µm width and 1.4 µm height and 25 µm length. The chips were made of polydimethysiloxane (PDMS) polymer using a silicon wafer mold (Conscience). A 1:10 solution of polymerising agent and PDMS monomer were rigorously mixed and then poured onto the silicon wafer. This was placed in a vacuum chamber and pressurised to remove air bubbles. The device was then heated at 65˚C in an oven for 2 hours to polymerise. For each experiment, one chip was cut out using a scalpel, and holes for inlet and outlet were inserted using a 0.75 mm biopsy puncher. The device was cleaned using 100% ethanol and dried with nitrogen gas. The cleaning was repeated 3 times. The PDMS chip was bonded on a glass coverslip (thickness No 1.5). These coverslips were first cleaned by sonication with acetone for 20 mins followed by isopropanol for 20 min, and then dried with nitrogen gas. The cleaned coverslip and PDMS chip were exposed to air plasma for 2 min and bonded at 95˚C for 30 min.

#### Mother machine setup

1 mL of exponentially grown cells were spun down for 2 min at 6000 rpm. These cells were then resuspended in 100 µL of the supernatant and loaded in the microfluidic chips by pipetting through the inlet. For experiments with a lower and variable number of cells per trench, the cells were centrifuged and resuspended in 500 µL of supernatant before being loaded into the chip. The chip was then inserted into a custom-built centrifuge holder and spun at 5000 rpm for 10 min to aid the loading of cells into the growth trenches. 50 mL syringes were filled with M9 minimal media containing Pluronic F127 and $H_2O_2$ as indicated. The syringes were attached to silicon tubing (Tygon) and loaded onto syringe pumps (NewEra SyringePumpPro) to deliver media into chips at a constant flow rate of 2.5 mL per hour. Cells were initially grown without $H_2O_2$ for ~3 hours before switching the inlet media to a syringe containing $H_2O_2$ using a Y-junction attached to the inlet of the chip.

#### Time-lapse microscopy

Time lapse imaging was performed using a Nikon Ti-E inverted fluorescence microscope equipped with 100x NA 1.40 immersion oil objective, motorized stage, sCMOS camera (Hamamatsu Flash 4), LED excitation source (Lumencor SpectraX), and operated with a perfect focus system. Exposure times were 100 ms for $P_{RNAI}$-mKate2 ($\lambda$ = 555 nm), 75 ms for CFP reporters ($\lambda$ = 440 nm) and 300 ms for MutL-mYPet ($\lambda$ = 508 nm) using 50% of maximal LED excitation intensities. The excitation and emission lights were separated using a triband dichroic and individual emission filters. The microscope chamber (Okolabs) was maintained at 37˚C throughout the experiments. Images were captured every 3 min for the 3 emission channels. On average, 45 to 50 fields of view (FOV) were captured per experiment with each FOV containing 26 growth trenches. This yielded ~1200 independent growth trenches each containing a single mother cell and its progeny. With 8 cells per growth trench, on average ~10,000 cells were captured in total per time point. For cross correlation analysis experiments, we imaged $P_{RNAI}$-mKate2 for cell segmentation at 50 ms and CFP channel at 50 ms exposure times every 45 seconds. Since this was a shorter time duration between cycles, we could capture around 18 to 20 FOV which yielded ~500 cells per time point.

**Imaging of microcolonies**

Cells were streaked on LB plates with antibiotics (25 µg/mL kanamycin and 100 µg/mL ampicillin) and grown overnight at 37˚C. A colony was picked and grown overnight in 4 mL LB at 37˚C in a shaking culture. The next day, 2 µl spots of overnight culture were dropped on a LB agarose plate without letting the pipette tip touch the agar surface and grown for 2 hours at 37˚C. Using a scalpel, the agarose was cut surrounding a spot (1 cm by 1 cm). 8 µl of 10 mM $H_2O_2$ solution in LB was dropped close to the spot and let to dry for ~3 minutes. The agarose pad was then flipped onto a clean glass slide and then covered with a cap of a culture tube that was sealed with tape. Time lapse movies were performed on a Nikon Ti-E microscope equipped with a 100x NA 1.45 oil immersion objective, motorised stage, sCMOS camera (Photometrics Prime95B), LED excitation source (Lumencor SpectraX) and perfect focus system. Exposure times were 100 ms for $P_{RNAI}$-mKate2 ($\lambda$ = 555 nm), 75 ms for CFP reporters ($\lambda$ = 440 nm) and 300ms for MutL-mYPet ($\lambda$ = 508 nm) using 50% of maximal LED excitation intensities. The microscope chamber (Okolabs) was maintained at 37˚C throughout the experiments. Images were captured every 3 minutes for brightfield and the 3 fluorescence channels. On average, 10 to 20 fields of view (FOV) were recorded per experiment with each FOV capturing a section of the microcolony.

**Mother machine data processing and analysis**

Time lapse microscopy data were saved as .nd2 files and visualized in Fiji. The data were processed using the BACMMAN plugin in Fiji as described in (Ollion et al., 2019) and further analysed using custom Python scripts. Graphs were generated using MATLAB. Images were first pre-processed by BACMMAN using the $P_{RNAI}$-mKate2 fluorescence channel to stack all individual growth trenches and correct for experimental drift in x-y coordinates and image rotation. The outlines of cells in the growth trenches were then jointly segmented and tracked over time based on the $P_{RNAI}$-mKate2 fluorescence signal. The traces were visually inspected and manually corrected for errors in segmentation or lineage tracing using the BACMMAN software. The CFP fluorescence of the different reporter plasmids was extracted by overlaying the cell masks from the $P_{RNAI}$-mKate2 channel onto the CFP channel and computing the mean intensity over the cell area. BACMMAN software was also used to detect foci of the MutL-mYPet reporter for DNA mismatches within the cell masks. BACMMAN generated output in 4 excel files containing cell growth characteristic, $P_{RNAI}$-mKate2 intensity data, CFP intensity data, and MutL-mYPet foci detections. These files were then further analysed using a custom python pipeline as described in the following.

**Cell parameter calculations**

**Cell length (L):** The cell length was computed from the maximum distance between the points on the cell masks.

**Cell area (A):** The cell area was computed from the total area covered by the cell mask.

**Estimated cell surface area to volume ratio:** This ratio was empirically calculated as: $\rho = \frac{24L\,(L^2+A)}{A(3L^2-A)}$

A cell is considered as a cylinder capped with 2 hemispheres with the total length as L and radius as r.

$$\frac{SA}{V} = \rho = \frac{2\pi r(L-2r) + 4\pi r^2}{\pi r^2(L-2r) + \frac{4\pi r^3}{3}}$$

Simplifying the above equation and approximating $A = 2rL$. Here A is the cell area output from BACMMAN.

$$\rho = \frac{24L\,(L^2+A)}{A(3L^2-A)}$$

**Elongation Rate**: The instantaneous elongation rate was calculated based on the log-difference in cell length between consecutive frames as $\frac{\log(L_t) - \log(L_{t-\Delta t})}{\Delta t}$. For calculating the elongation rates of cells at different

positions in the growth trench (e.g. Fig 1C), cells were tracked according to their initial position until the number of barrier cells decreased by 2.
**Generation time:** The time difference between two consecutive cell divisions.
**Division Rate**: The inverse of the cell generation time.
**Reporter fluorescence intensities**: The intensity values were averaged over the area of each cell.
**Length Growth Rate**: This was calculated as the coefficient of a linear fit (β) to the logarithm of cell length over each cell cycle. $L = L_0 e^{\beta t}$, where $L_0$ is the length at birth. Since, it is calculated over the whole lineage duration, we cannot capture the instantaneous changes in elongation rate.
**Fate of a cell**: A mother cell was defined dead if the length growth rate was less than 0.012 $min^{-1}$ for >10 hours after stress removal.
**Number of barrier cells:** The number of cells that are located between the open end of a trench and the cell being analysed.
**Cumulative barrier length**: The sum of lengths of barrier cells.
**Cumulative barrier area**: The total area covered by the barrier cells.
**Cumulative barrier surface area to volume**: The total estimated surface area to volume ratio of all barrier cells.
**Lineage annotation:** identity of cells was obtained from BACMMAN to define their lineage relations as sisters, cousins, or non-related.

**Stress response parameter calculations**

**Response peak intensity**: The oxidative stress response reporters showed an initial peak in intensity and then reached a lower steady-state level. The initial peak intensities for each mother cell were detected using the PeakUtils module in Python with a normalised threshold of 0.15 and minimum distance between peaks of 2 frames. This absolute response peak value was subtracted by the mean reporter intensity without $H_2O_2$ treatment.
**Response induction time:** The time at which a mother cell crosses an intensity threshold of 20% of the steady state intensity value (i.e. 1480 a.u. for P*grxA*-CFP) after the start of $H_2O_2$ treatment.
**Response peak time:** The time at which a mother cell reached its response peak intensity after the start of $H_2O_2$ treatment.
**Coefficient of variation**: The CV values were calculated as the mean divided by the standard deviation for the response induction time, peak time, and peak intensity. CVs of steady state intensities were calculated for mother cells or all cells from 90 minutes post treatment until 6 hours post treatment after subtracting the mean intensity for each cell before $H_2O_2$ treatment.

**Cross-correlation analysis**

The temporal cross-correlation between the reporter intensity traces of mother cells and the number of barrier cells per trench was computed using the statsmodel library in Python. Correlation values from individual growth trenches were then averaged over all observed growth trenches for different lag times. The minimum of the cross-correlation curve was obtained using the argrelextrema function in Python to obtain the lag time of the reporter intensity in response to changes in the number of barrier cells.

**Calibration of $H_2O_2$ concentration inside growth trenches**

We estimated the local $H_2O_2$ concentration in the growth trenches from the P*grxA*-CFP reporter intensities of cells located at different positions. We calibrated the analysis using the reporter intensities of the cells located at the open end of the growth trenches that were exposed to defined $H_2O_2$ concentrations (12.5 μM, 25 μM, 37.5 μM, 50 μM, 62.5 μM, 75 μM and 100 μM). The fluorescence intensities at steady-state (1- 3 hours after start of treatment) showed a linear relation with the external $H_2O_2$ concentration. This was used to obtain the conversion factor for computing the $H_2O_2$ concentration from the reporter intensity of cells at any position in the trenches.

**Machine learning**

We trained a random forest regressor model to predict the initial peak intensity of the P*grxA*-CFP reporter for each mother cell according to the schematic in Figure 2A. A separate random forest classifier model was trained to predict the external $H_2O_2$ concentration. A list of features (shown below) was calculated from the output data provided by BACMMAN. Data from multiple experiments (as indicated in figure captions) were combined into one dataset for model training and testing. Feature values for each mother cell and its environment were stored in separate CSV files which served as input to the Python-based Machine learning model. The following Python libraries/packages were used: pandas; numpy; peakutils; matplotlib; stats, from scipy; preprocessing, utils, shuffle, metrics, model_selection, ensemble from sklearn, ststamodel.api.

126 features were extracted from each mother cell and labelled with the output value i.e. the P*grxA*-CFP peak intensity for each cell. The data was then split 80:20 into training and test data using train_test_split module from sklearn. The training and test data were then individually shuffled using the shuffle library in sklearn. Utilising the supervised learning method, a random forest model is an ensemble technique that can predict the output numerical value or classification class, for regressor and classifier respectively, based on the combined outputs generated by multiple decision trees that each process a random subset of the data with a random subset of the available features (Breiman, 2001).
For the regression model, the decision trees compute numerical values at each tree node. The feature values were extracted from a subset of training data at each decision node that were randomly drawn from the feature list. Prediction was then performed for multiple trees and mean squared error was calculated for every predictor and minimized by bootstrapping. The final output numerical value for the P*grxA*-CFP peak intensity per cell is the average of these predictions.
The classification model generates probabilistic predictions of labelled outputs of the given data set, whose decision is based on bootstrapping results from a collection of randomized trees. Here, the different $H_2O_2$ treatment concentrations were considered as separate classes for each mother cell. The decisions trees making up the forest were trained with the input data as an independent classifier. These independent trees are formed using random subsets from training data sets and the randomized feature set that was extracted. The label that is selected by the majority of the trees is output as the class.

We used 100 trees and forest depth of 3000 for bootstrap samples in both of the above-mentioned algorithms. After the models had been trained, the test data were used to evaluate the model accuracy based on the normalized difference between the predicted and observed P*grxA*-CFP peak intensity per cell, and averaged over all cells in the test dataset. We report the model accuracy accordingly:

$$A = 1 - \frac{1}{n}.\sum_{s=1}^{s=n} \left| \frac{pred_s - exp_s}{exp_s} \right|$$

The accuracy of the classification model was evaluated using a confusion matrix to quantify the frequency of true positive and true negative predictions. To assess the relative importance of the different feature types for the predictive power of the model, we used the mean decrease impurity test in the scikit-learn package in Python (Breiman, 2001).

### Machine Learning Features

$$GrxA_{peak} = \{GrxA|_{t_p<t<t_q}\}_{peak} - \frac{\sum_{t=-treatment}^{t=0} GrxA}{treatment} \; ; t_p = 0 \, , t_q = 60 \; minutes$$

1. Kurtosis = $Y_{kurt} = (t_2 - t_1) * \frac{\sum_{t=t_1}^{t=t_2}\left(Y_t - \frac{\sum_{t=t_1}^{t=t_2} Y_t}{t_2 - t_1}\right)}{\left(\sum_{t=t_1}^{t=t_2}\left(\left(Y_t - \frac{\sum_{t=t_1}^{t=t_2} Y_t}{t_2 - t_1}\right)^2\right)\right)^2}$
2. Skewness = $Y_{skew} = (t_2 - t_1) * \frac{\frac{\sum_{t=t_1}^{t=t_2} Y_t}{t_2 - t_1} - Median(Y_t \,\forall\, t_1 < t < t_2)}{\left(\frac{\sum_{t=t_1}^{t=t_2}\left(Y_t - \frac{\sum_{t=t_1}^{t=t_2} Y_t}{t_2 - t_1}\right)^2}{t_2 - t_1}\right)^{0.5}}$
3. Median = $Y_{median} = Median(Y_t \,\forall\, t_1 < t < t_2)$
4. Mean or average = $Y_{mean} = \frac{\sum_{t=t_1}^{t=t_2} Y_t}{t_2 - t_1}$
5. Minimum = $Y_{min} = Minima(Y_t \,\forall\, t_1 < t < t_2)$
6. Maximum = $Y_{max} = Maxima(Y_t \,\forall\, t_1 < t < t_2)$
7. Range of magnitude = $Y_{range} = Y_{max} - Y_{min}$

**All the above features were calculated for 'mother cells' where $Y_t$ =:**

1. Cell length at time point t = $l_t$
2. Cell area at time point t = $A_t$

3. Instantaneous cell elongation rate at time point t = $\beta_t = \frac{l_t - l_{t-1}}{1}$
4. Instantaneous cell area increase rate at time point t = $\rho_t = \frac{A_t - A_{t-1}}{1}$
5. Estimated cell surface area to volume ratio at time point t = $\alpha_t = \frac{l_t}{A_t - (\frac{2A_t^2}{3l_t^2})}$
6. Number of barrier cells at time point t =$B_t = Maxima(Idx_{cell,t}) - 1$
7. Total length of barrier cells =$l_{t,barrier} = (\sum_{i=0}^{i=B_t} l_i)_t$
8. Total area of barrier cells = $A_{t,barrier} = (\sum_{i=0}^{i=B_t} A_i)_t$
9. Cumulative 3D surface area to volume ratio of barrier cells = $\alpha_{t,barrier} = \left( \sum_{i=0}^{i=B_t} \frac{l_i}{A_i - \left(\frac{2A_i^2}{3l_i^2}\right)} \right)_t$

**For accounting for different conditions, the following time-point series were analysed:**

1. Untreated : $t_2 = \text{treatment}, t_1 = \text{treatment} - 135 \text{ minutes}$
2. Hydrogen peroxide treated : $t_2 = \text{treatment} + 135 \text{ minutes}, t_1 = \text{treatment}$

Total number of input features = 7*9*2 = 126 + 1 output variable = 127 features

**Microcolony image analysis**

Segmentation of cells growing in microcolonies was performed based on the $P_{RNAI}$-mKate2 fluorescence signal and using the MicrobeTracker tool in MATLAB (Sliusarenko et al., 2011) followed by manual correction of the segmentation masks. These outlines were then applied to the CFP channel and a MATLAB script was used to quantify the average intensity per cell area.

**MutL-mYPet foci detection**

MutL-mYPet foci detection was carried out by BACMMAN. Images were band pass filtered between within 1 to 400 pixels. Next, the image was gaussian smoothed with a scale of 2 pixels and the spot segmentation was performed using Seed Laplacian threshold of 0.8, propagation threshold of 1.5 and seed threshold of 1.7. Foci that were detected in multiple consecutive frames within the same cell were counted only once in the first frame. The mismatch rate was calculated by dividing the total number of foci by the total number of cells per frame.

| REAGENT or RESOURCE | SOURCE | IDENTIFIER |
|---|---|---|
| Bacterial Strains | | |
| AB1157, Δ*flhD*, $P_{RNAI}$-mKate2, mutL-mYpet (SU178) | (Lagage et al., 2022) | N/A |
| AB1157, Δ*flhD*, $P_{RNAI}$-mKate2, mutL-mYPet, carrying pUA139 P*grxA*-SCFP3A Kan (SU777) | (Lagage et al., 2022) | N/A |
| AB1157, Δ*flhD*, $P_{RNAI}$-mKate2, mutL-mYpet, Δ*oxyR*, carrying pUA139 P*grxA*-SCFP3A kan (PSU044) (SU802) | (Lagage et al., 2022) | N/A |
| AB1157, Δ*flhD*, $P_{RNAI}$-mKate2, mutL-mYpet, carrying pUA066-P*katG*-SCFP3A kan (SU620) | (Lagage et al., 2022) | N/A |
| AB1157, Δ*flhD*, $P_{RNAI}$-mKate2, mutL-mYpet, carrying pUA066-P*ahpC*-SCFP3A kan (SU948) | This study | N/A |
| AB1157, Δ*flhD*, $P_{RNAI}$-mKate2, MutL-mYPet carrying P*katE*-SCFP3A kan (SU945) | This study | N/A |
| AB1157, Δ*flhD*, $P_{RNAI}$-mKate2 P*grxA*-SCFP3A kan (SU880) | This study | N/A |

| | | |
|---|---|---|
| AB1157, Δ*flhD*, mKate2, MutL-mYPet, Δ*oxyR,* pUA P*grxA*-SCFP3A kan + pUC18 amp encodes YPet preceeded by 11 aa linker followed by a kan cassette flanked by frt sites. (SU882) | This study | N/A |
| Chemicals, Peptides, and Recombinant Proteins | | |
| M9 minimal salts 5x | Sigma | M9956 |
| MEM amino acids | Gibco | 11130-036 |
| L-Proline | Biochemica | A3453,0100 |
| Thiamine | Biochemica | A0955,0050 |
| Pluronic F-127 | Sigma | P2443-250G |
| 30% W/W solution of H2O2 | Sigma | H1009-100mL |
| Software and Algorithms | | |
| MATLAB | Mathworks | Mathworks.com |
| BACMMAN | Fiji | (Ollion et al., 2019) |
| Python | Spyder | anaconda.com |

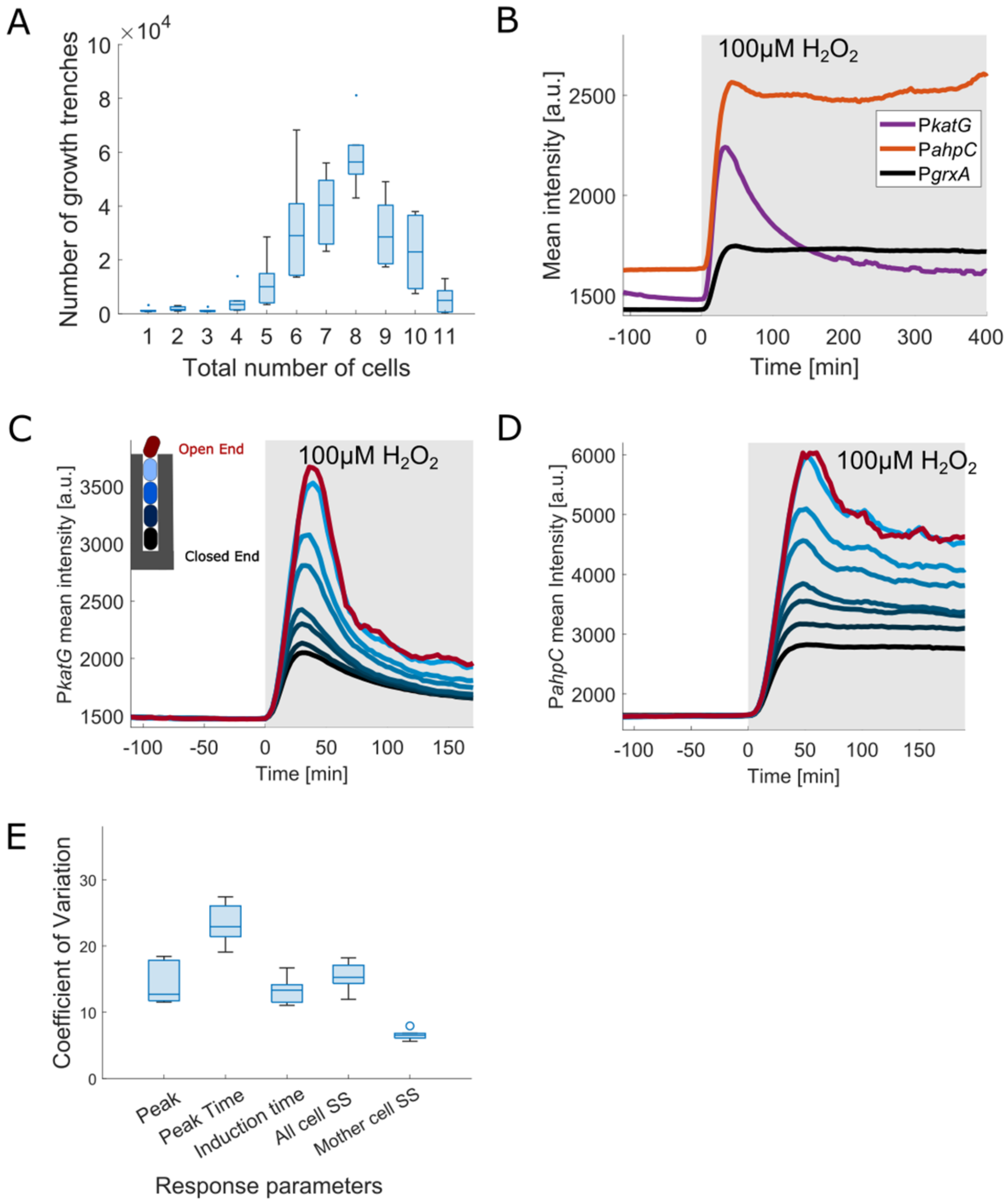


Figure S1: **Oxidative stress response characterised in microfluidic experiment:** (A) Distribution of number of cells per growth trench at the time of 100 µM $H_2O_2$ treatment (3 experimental repeats). (B) Mean CFP intensity for mother cells under 100 µM $H_2O_2$ treatment added at time 0 min (shaded area) for the transcriptional reporters of P*katG* and P*ahpC* and P*grxA* (≥ 2 experimental repeats each). (C) P*katG*-CFP intensities with continuous 100 µM $H_2O_2$ treatment added at time 0 min (shaded area) averaged across cells at specific positions in the growth trench (2 experimental repeats, black line: mother cells at closed end; red line: cells at open end). (D) P*ahpC*-CFP plot similar to panel E (2 experimental repeats). (E) Variation of the oxidative stress response across mother cells with 100 µM $H_2O_2$ treatment. Coefficient of Variation (standard deviation/mean) for the peak amplitude (Peak), the time to reach the PgrxA-CFP peak intensity (Peak Time), the response induction time (time until P*grxA*-CFP > 1480 a.u.) and P*grxA*-CFP intensity from 2 hours post treatment (SS: steady-state) for all cells in growth trenches and for all mother cells (3 experimental repeats, box plots with median $25^{th}$ and $75^{th}$ percentile).

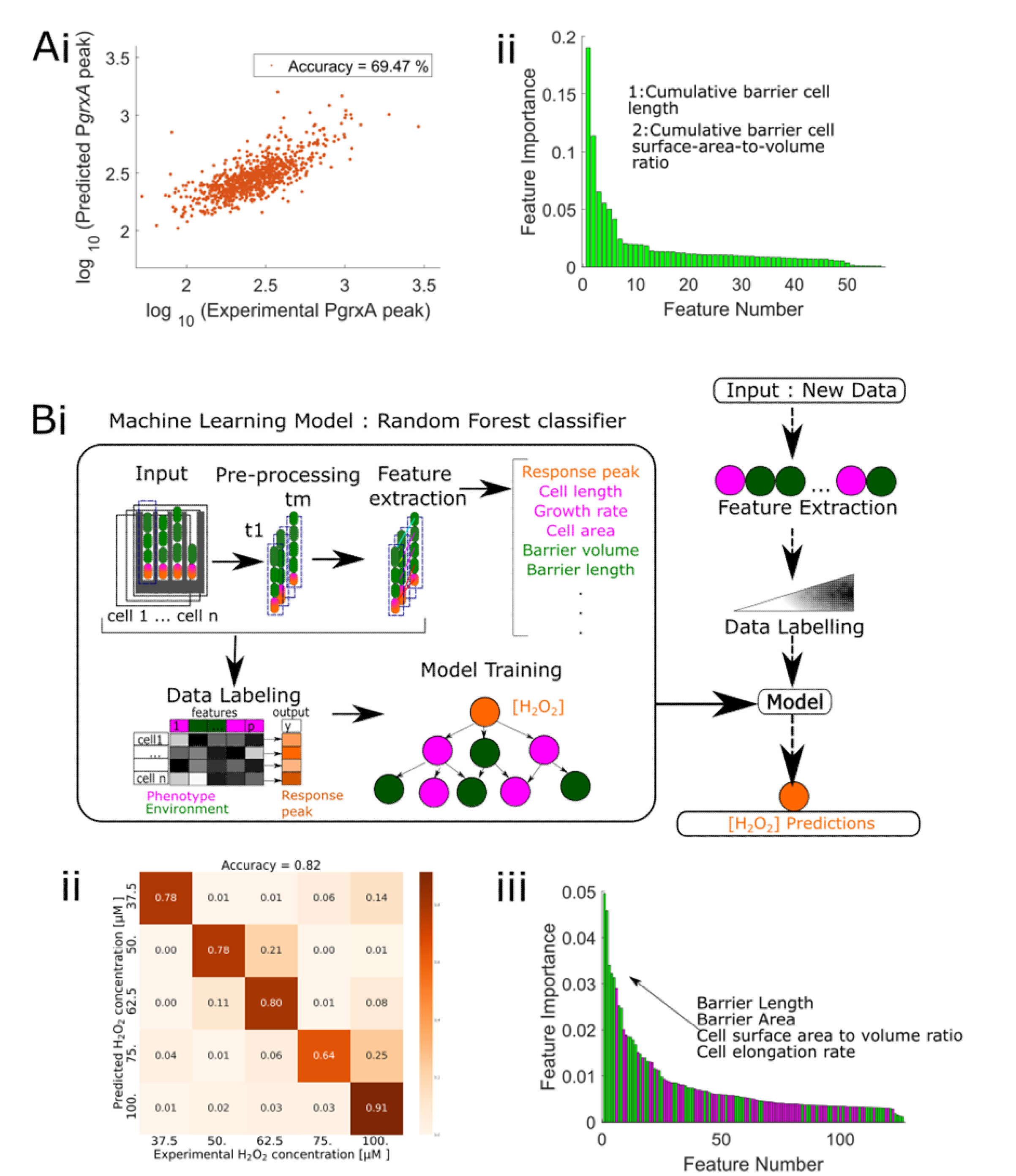


Figure S2: **Machine learning models to predict oxidative stress response heterogeneity and $H_2O_2$ treatment concentrations.** (A) Machine learning model trained only on barrier cells predicts mother cell response heterogeneity: A random forest machine learning model predicts P*grxA*-CFP peak intensities of ~850 mother cells (orange). It uses features that describe the other cells in the local environment of each trench (barrier cells, green), and no features for the mother cells themselves. (i) P*grxA*-CFP peak predicted by that model plotted against the experimentally measured P*grxA*-CFP peak (each dot represents one mother cell) (data in Table S6). (ii) Feature importance plot shows the relative contribution of the 54 input features to the predictive power of the model. The features of the two most important environmental characteristics accounting for ~73.5% importance are highlighted (feature names in Table S5). (B) Machine learning classifier model predicts external $H_2O_2$ concentration(orange). (i) It uses features that describe the phenotypic characteristics of the mother cell (magenta) and the other cells in the local environment of each trench (barrier cells, green). (ii) Confusion matrix of predicted [$H_2O_2$] against the experimental [$H_2O_2$] for ~900 mother cells (unseen by training data) (data shown in Table S4). (iii) Feature importance plot shows the relative contribution of the 126 input features to the predictive power of the model. Mother cell features shown in magenta and local environment features (relating to the barrier cells) in green. The features whose mathematical derivatives are in the top 10 most important features are highlighted. Whereas the prediction of P*grxA*-CFP peak intensities relied on only a few important features relating to the barrier cells, prediction of [$H_2O_2$] uses a broader range of different features relating to the mother cell and barrier cells (feature names in Table S3).

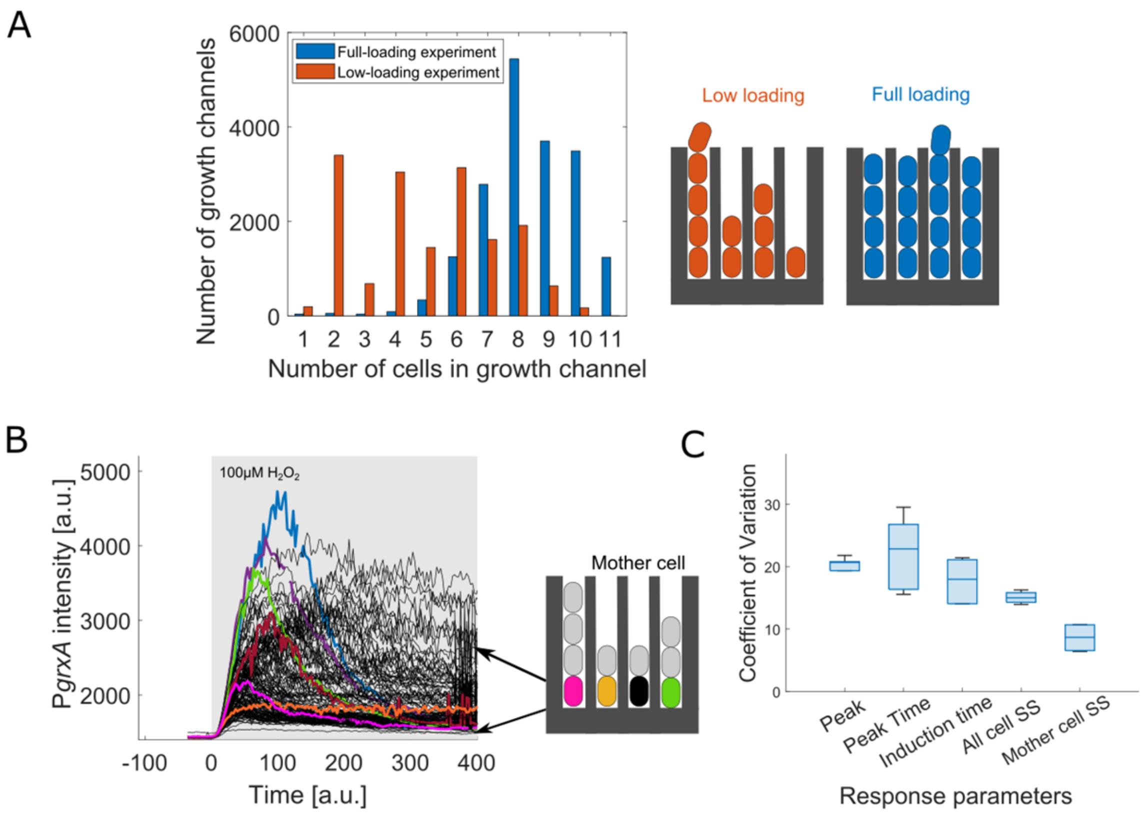


Figure S3: **Variation in the number of barrier cells per growth trench increases response heterogeneity across mother cells**: (A) Distribution of number of cells per trench at the time of treatment for experiments with partially filled trenches (low-loading, orange) versus completely filled trenches (full-loading, blue) (3 experimental repeats each). (B) P*grxA*-CFP intensities of individual mother cells (6 example cells highlighted in colour) over time treated with 100 µM $H_2O_2$ (added at time 0 min, shaded area) (~300 traces). (C) Variation of the oxidative stress response across mother cells for low-loading experiment after 100 µM $H_2O_2$ treatment. Coefficient of Variation (standard deviation/mean) for the peak amplitude (Peak), the time to reach the PgrxA-CFP peak intensity (Peak Time), the response induction time (time until P*grxA*-CFP > 1480 a.u.) and P*grxA*-CFP intensity from 2 hours post treatment (SS i.e. steady state) for all cells in growth trenches and for all mother cells (3 experimental repeats, box plots with median 25$^{th}$ and 75$^{th}$ percentile).

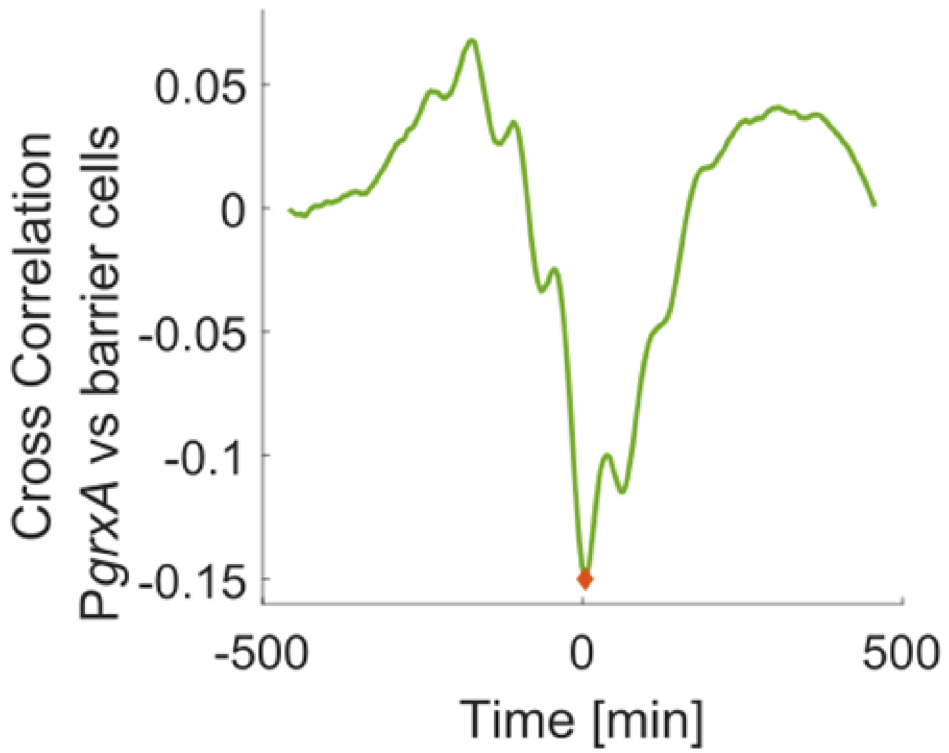


Figure S4: **Response fluctuations of mother cells are negatively correlated with variation in the number of barrier cells at steady-state.** Mean temporal cross correlation for P*grxA*-CFP of mother cells against the number of barrier cells per trench (example time traces shown in Figure 3c), when mean P*grxA*-CFP intensity has reached steady-state from 2 hours after start of 100 µM $H_2O_2$ treatment until end of experiment (~11 hours) (~950 cells, 2 experimental repeats). Measurements were performed with 45 second time interval between frames.

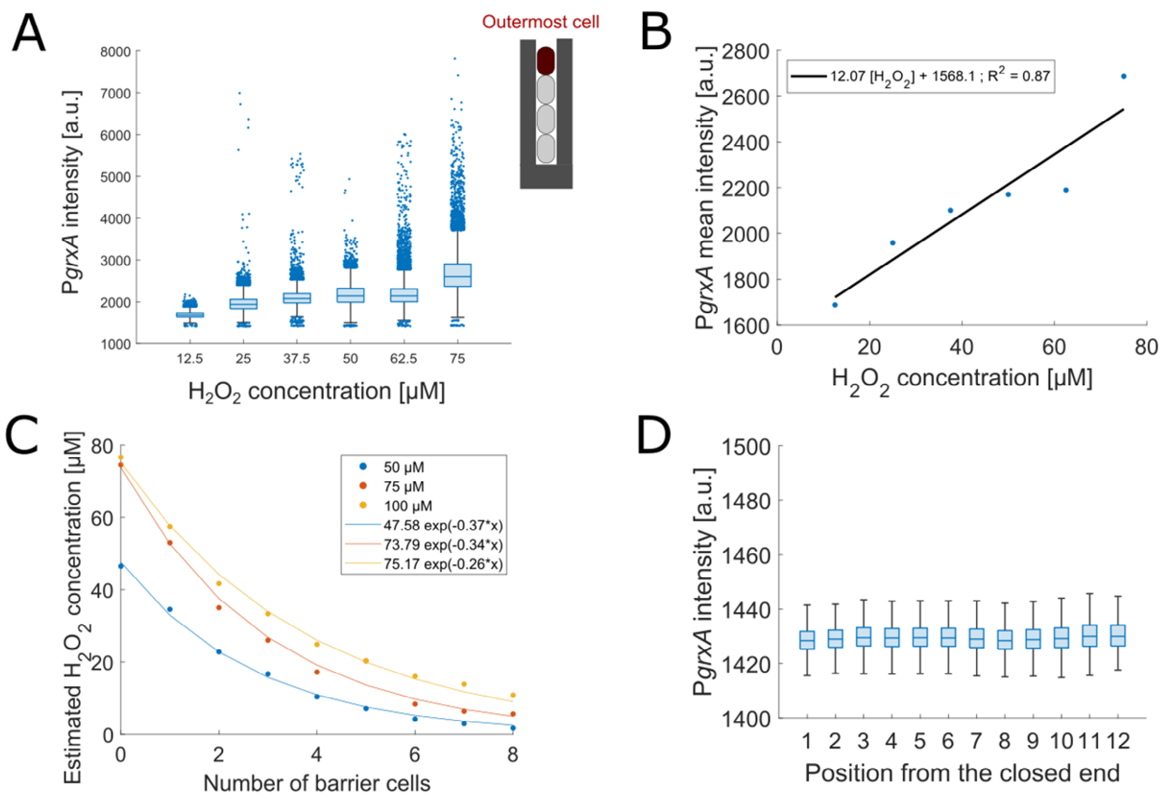


Figure S5: **Exponential decay of $H_2O_2$ in growth trenches as predicted by the calibration curve**: (A) P*grxA*-CFP intensities for these outermost cells 2 hours after the start of treatment for different $H_2O_2$ concentrations. (~107500 data points with 17921 ± 1960 data-points for each concentration, box plots with median 25th and 75th percentile) (B) A linear regression fit of the mean values in panel A gave the calibration equation for intensity changes for different $H_2O_2$ concentrations as $I = 12.07 \cdot [H_2O_2] + 1568.1$ with $R^2$=0.87. (C) Estimated concentration along the growth trench with mean intensities for varying number of barrier cells based on the PgrxA-CFP intensity according to the calibration curve from panel B. Single exponential fits are shown. (D) P*grxA*-CFP intensities for cells at different positions in the growth trench before treatment (3 experimental repeats, box plots with median 25th and 75th percentile).

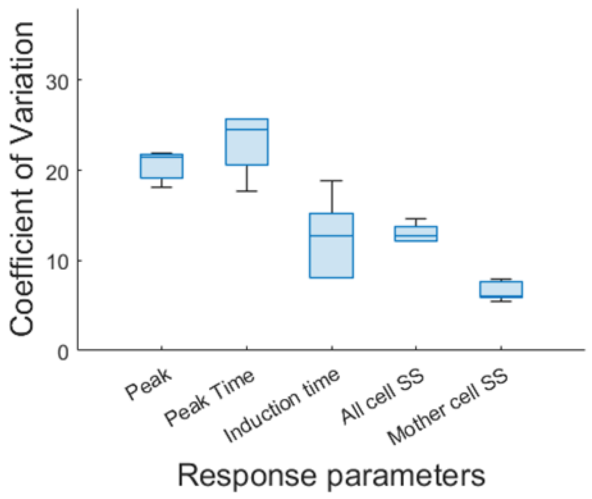


Figure S6: **Variation of the oxidative stress response across mother cells in 1.4 µm wide growth trenches.** Variation of the oxidative stress response across mother cells with 100 µM $H_2O_2$ treatment. Coefficient of Variation (standard deviation/mean) for the peak amplitude (Peak), the time to reach the P*grxA*-CFP peak intensity (Peak Time), the response induction time (time until P*grxA*-CFP > 1480 a.u.) and P*grxA*-CFP intensity from 2 hours post treatment (SS: steady-state) for all cells in growth trenches and for all mother cells (3 experimental repeats, box plots with median 25$^{th}$ and 75$^{th}$ percentile).

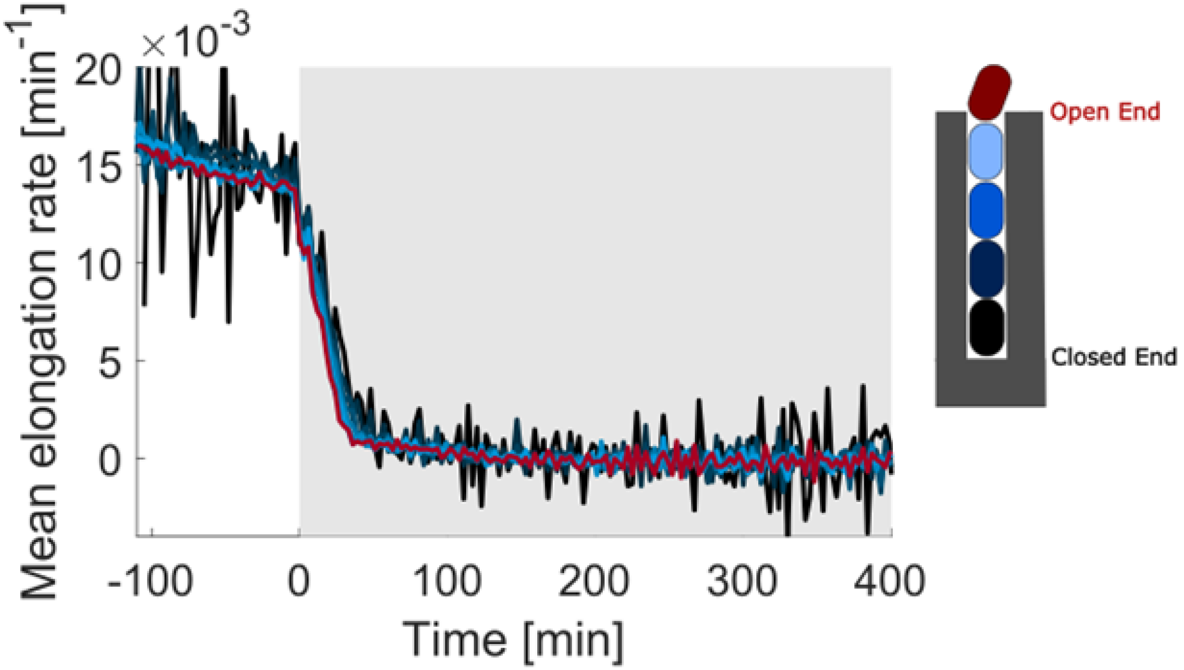


Figure S7: **Cells with Δ*oxyR* deletion do not survive 100 μM $H_2O_2$:** Mean elongation rate for Δ*oxyR* cells at different positions in the growth trench with 100 μM $H_2O_2$ treatment (shaded area; black line: mother cells at closed end; red line: cells at open end) (3 experimental repeats).

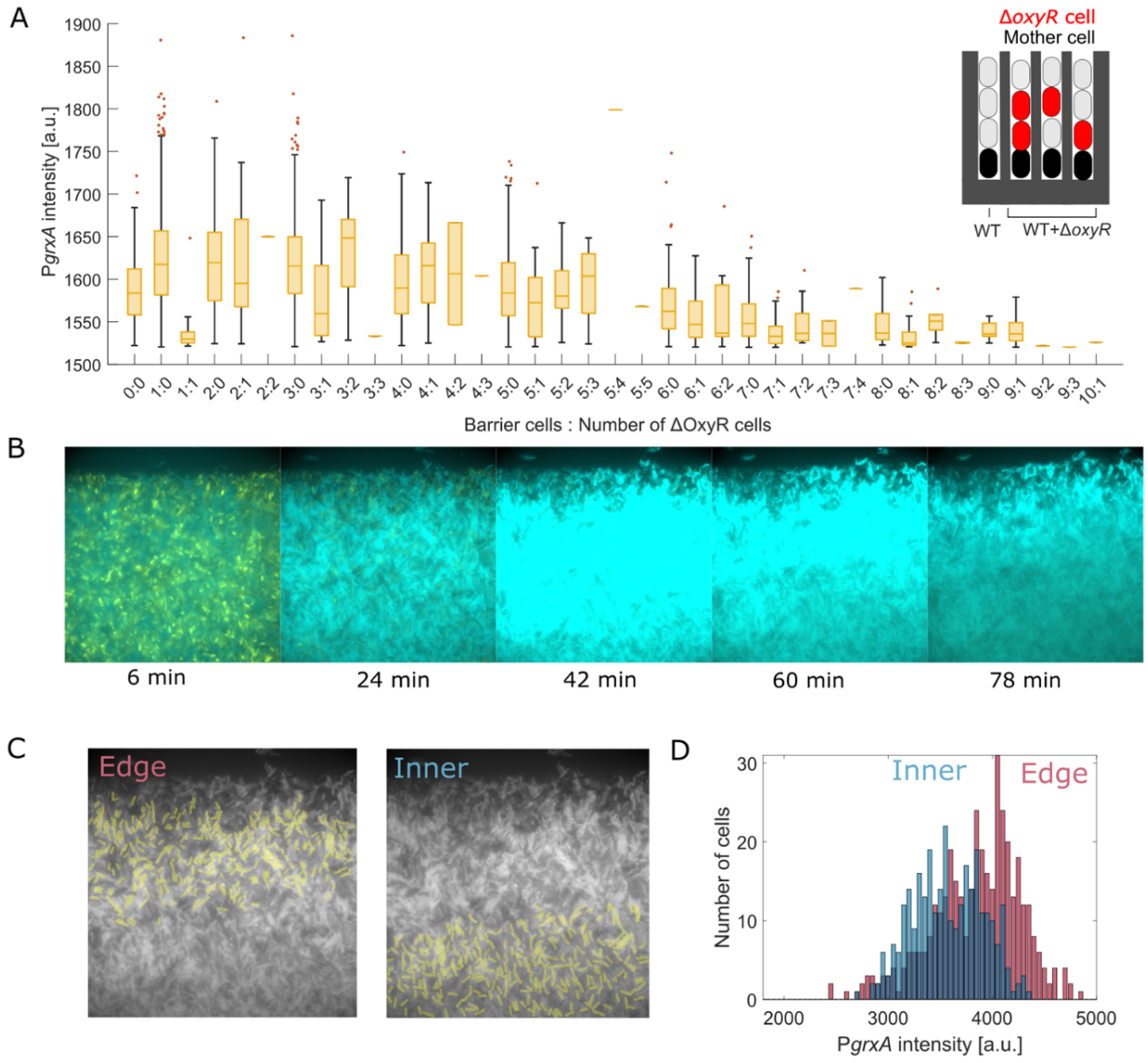


Figure S8: **Δ*oxyR* cells do not provide cross protection against $H_2O_2$ stress**: (A) P*grxA*-CFP intensity 30 min post 100 μM $H_2O_2$ treatment for WT mother cells in trenches that have variable mixture of WT and Δ*oxyR* cells. The ratio of total number of barrier cells to Δ*oxyR* cells per trench is shown on the x-axis (3 experimental repeats, box plots with median 25$^{th}$ and 75$^{th}$ percentile). (B) P*grxA*-CFP snapshots of a microcolony of WT mixed with Δ*oxyR* cells (yellow) under 10 mM $H_2O_2$ treatment. (C) Segmented WT cells on the edge or interior of the microcolony. (D) Histograms of P*grxA*-CFP intensity for WT cells at the edge or interior of a microcolony after 30 min of 10 mM $H_2O_2$ treatment.

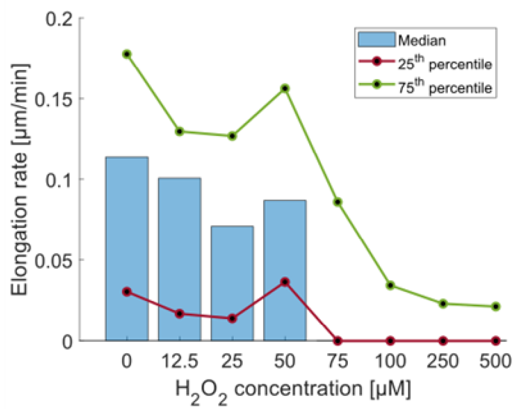


Figure S9: **Growth inhibition in the absence of cellular cross-protection.** Elongation rate for outermost cells in the trenches traced from time of treatment until 30 minutes post treatment for different $H_2O_2$ concentrations.

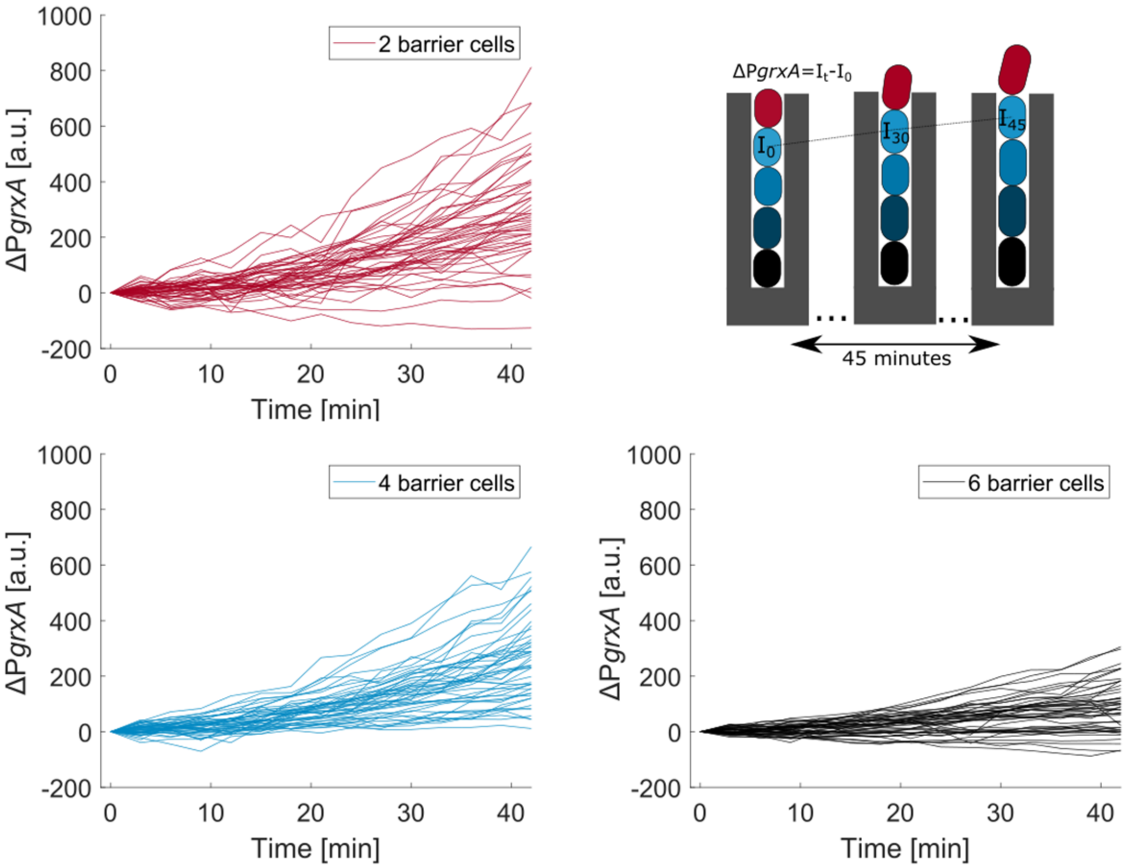


Figure S10**: Effect of cell movement on P*grxA*-CFP expression in a spatial $H_2O_2$ gradient.** Increase in P*grxA*-CFP intensity for single cells (Δ*PgrxA*) over time when the response has reached steady-state (from 2 hours after start of 100 µM $H_2O_2$ treatment until end of experiment ~11 hours). Each curve represents a single cell moving towards the trench opening from a different starting position. (red line: cells with 2 barriers; blue line: cells with 4 barriers; black line: cells with 6 barriers; 45 traces shown for each case).

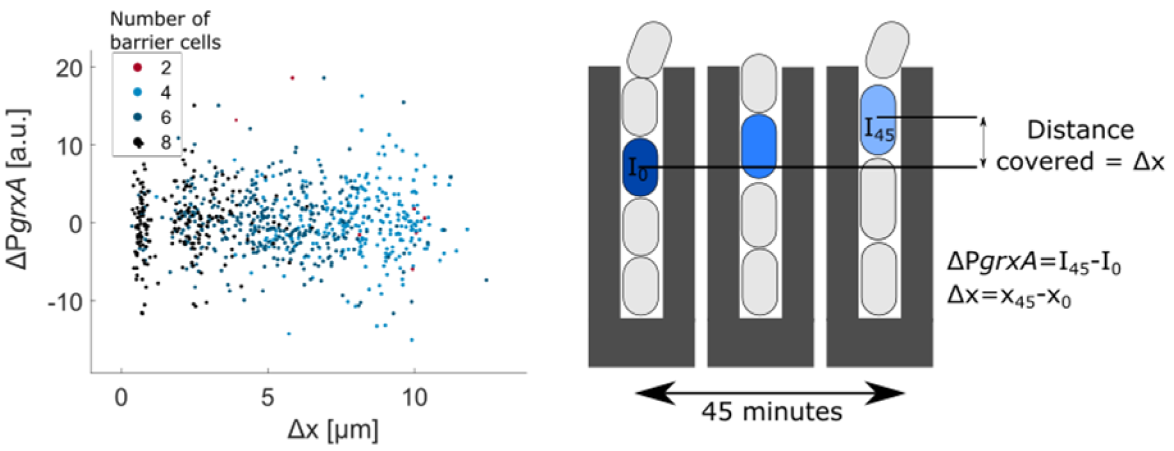


Figure S11**: Cell movement in trenches does not affect P*grxA*-CFP intensity in untreated conditions:** Each point represents the total increase in P*grxA*-CFP and total increase in distance for single cell traced for 45 minutes without $H_2O_2$ treatment (black line: closer to the closed end; red line: cells at open end) (~800 cells, 3 experimental repeats).

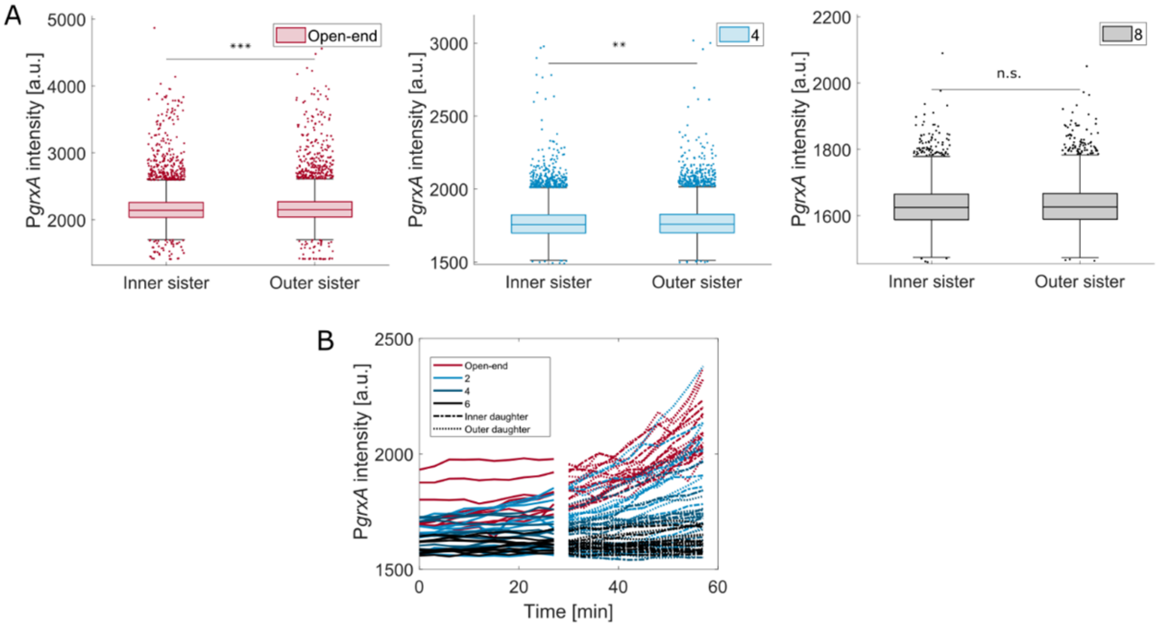


Figure S12: **Response of sister cells diverge under 100 µM $H_2O_2$ treatment**: (A) P*grxA*-CFP intensity at the time of division for sister cells at steady state with 100 µM $H_2O_2$ treatment. (Outer sister: closer to the open end). Pairs with outer sister having 0 (maroon), 4 (blue) and 8 (black) barrier cells are displayed (~ 34000 sister cell pairs, 3 experimental repeats, box plots with median 25$^{th}$ and 75$^{th}$ percentile; *** $p<0.001$, ** $p<0.01$ and n.s. for $p>0.05$). (B) Single-cell traces for the data shown in Figure 6F. P*grxA*-CFP intensity traces for sister cells (outer: dashed, inner: dash-dotted), and their progenitor cells (time of division ± 27 minutes). Colour code based on number of barrier cells at the time of division (10 mother-daughter pairs for each case plotted).

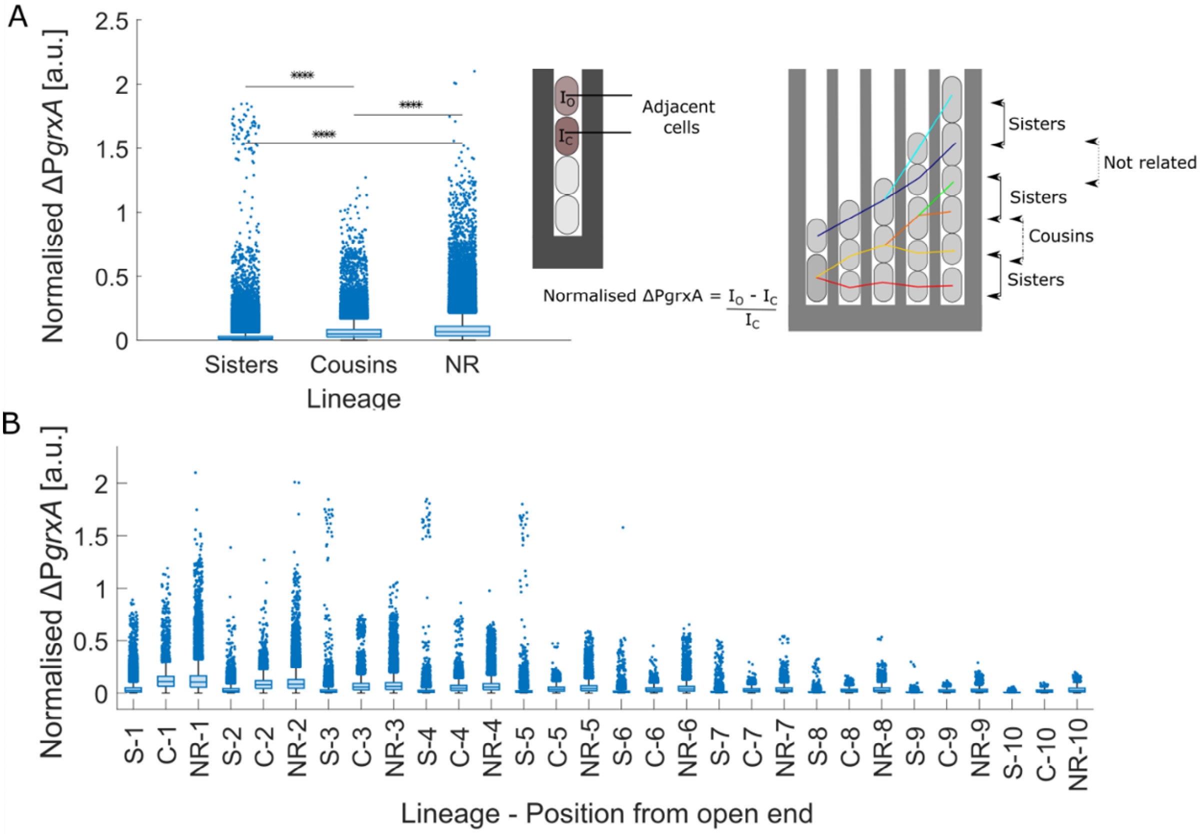


Figure S13: **Cells become more correlated with closer lineage relation and increasing barrier lengths**: (A) Normalised P*grxA*-CFP intensity difference for adjacent cell pairs related as sister (S), cousins (C), or not related (NR) under 100 µM $H_2O_2$ (box plots with median 25$^{th}$ and 75$^{th}$ percentile, **** p<0.0001, 3 experimental repeats). (B) Normalised P*grxA*-CFP intensity differences for adjacent cells at different positions from the open end and separated by lineage identity (e.g. S-1 shows the intensity difference between sisters at position 1 from the open end etc; 3 experimental repeats, box plots with median 25$^{th}$ and 75$^{th}$ percentile). In general, unrelated cells have higher intensity differences than cousins or sisters, and intensity differences decrease with increasing number of barrier cells.

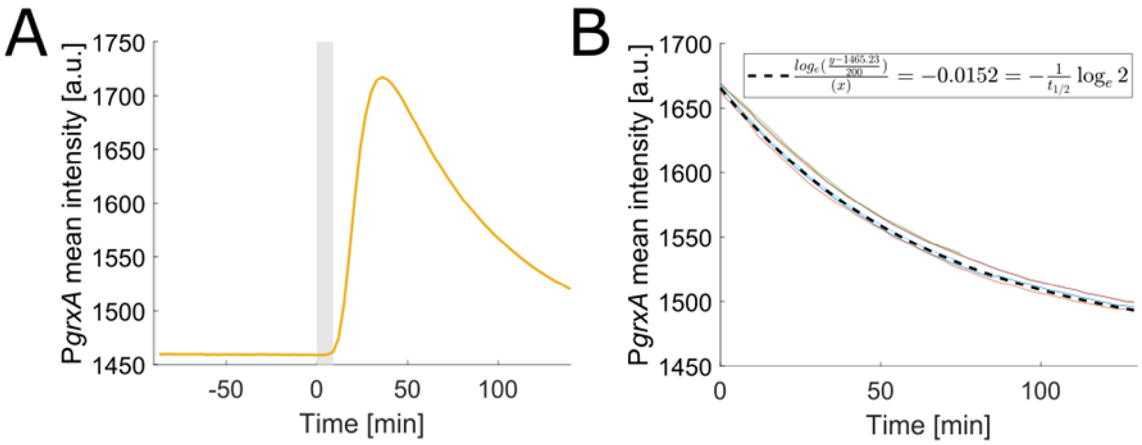


Figure S14: **Exponential response decay after $H_2O_2$ removal:** (A) Mean P*grxA*-CFP intensity for 100 µM $H_2O_2$ treatment from 0 to 9 minutes (shaded). (B) Exponential fit to the decay of intensity after treatment removal of data shown in panel A, $t_{1/2}$ = 45.6 min as expected for the generation time of ~45 min (thin lines show curve of different dilution traces and bold dashed line indicates the fit, 6 repeats). This shows that sCFP3 is stable and diluted by cell growth.

**Movie S1**: P*grxA*-SCFP3 fluorescence of WT cells with 100 μM $H_2O_2$ treatment from time 0. Related to Figure 1.

**Movie S2**: Mix of WT and Δ*oxyR* cells under 100 μM $H_2O_2$ treatment. Related to Figure 4. Treatment was provided at time = 0. The RFP channel overlaid with CFP and YFP is shown in the movie to visualise cell boundary with $P_{RNAI}$-mKate2 expression, P*grxA*-SCFP3 expression and DNA mismatch detected as MutL-mYPet foci.

**Movie S3**: Δ*oxyR* cells under 100 μM $H_2O_2$ treatment. Related to Figure S7. Treatment was provided at time = 0. The RFP channel overlaid with YFP is shown in the movie to visualise cell boundary with the $P_{RNAI}$-mKate2 expression and DNA mismatch detected as MutL-mYPet foci.

**Movie S4:** The presence of wild-type cells acting as a barrier rescues the growth of Δ*oxyR* cells until the wild-type cells exited the trench. Mix of WT and Δ*oxyR* cells in growth trenches 1, 2, 4 and 7 cells with Δ*oxyR* cells occupying mother cell position in trenches 1, 2 and 4. Treatment was provided at time = 0. The RFP channel overlaid with CFP and YFP is shown in the movie to visualise cell boundary with $P_{RNAI}$-mKate2 expression, P*grxA*-SCFP3 expression and DNA mismatch detected as MutL-mYPet foci.

**Movie S5:** The presence of wild-type cells acting as a barrier rescues the growth of Δ*oxyR* cells until the wild-type cells exited the trench. Mix of WT and Δ*oxyR* cells in the right growth trench with Δ*oxyR* cell occupying the mother cell position. Treatment was provided at time = 0. The RFP channel overlaid with CFP and YFP is shown in the movie to visualise cell boundary with $P_{RNAI}$-mKate2 expression, P*grxA*-SCFP3 expression and DNA mismatch detected as MutL-mYPet foci.

**Movie S6**: WT colony under 10 mM $H_2O_2$ treatment. Images were taken every 3 minutes, starting 6 minutes post treatment. The YFP channel showing MutL-mYPet foci overlaid with P*grxA*-SCFP3 is shown.

**Table S1**: Machine learning regressor model feature importance list. Related to Figure 2C. The description for all features is in the material and methods section. [RFR_featureList.csv]

**Table S2**: Predicted and experimental P*grxA* peak values for test data output by the regressor model. [RFR_error.csv]

**Table S3**: Machine learning classifier model feature importance list. Related to Figure S2B. The description for all features is in the material and methods section. [RFC_featureList.csv]

**Table S4**: Predicted and experimental $H_2O_2$ concentration values for test data output by the classification model. [RFC_error.csv]

**Table S5**: Machine learning regressor model feature importance list for model trained only on barrier cell features. Related to Figure S2A. The description for all features is in the material and methods section. [RFR_featureList_onlybarrier.csv]

**Table S6**: Predicted and experimental P*grxA* peak values for test data output by the regressor model trained only on barrier cell features. [RFR_error_onlybarrier.csv]
